# A modular generalist-specialist AI framework for ROI selection across spatial profiling workflow

**DOI:** 10.64898/2026.06.26.734862

**Authors:** Simon P. Castillo, Tanishq Gautam, Karina B. Pinao Gonzales, Maria E. Salvatierra, Alejandra Serrano, Caner Ercan, B. Leticia Rodriguez, Paul Acosta, Pingjun Chen, Yasin Shokrollahi, Alexandria Lau, Lawrence N. Kwong, Jason T. Huse, Xiaoxi Pan, Patient Mosaic Team, Luisa Solis-Soto, Yinyin Yuan

**Author notes:** Correspondence and requests for materials should be addressed to S.P.C. and Y.Y. Samantha Dickson Brain Cancer Unit, UCL Cancer Institute, University College London, London, WC1E 6DD, United Kingdom. Full collaborator list in Supplementary Material. These authors contributed equally: Simon P. Castillo, Tanishq Gautam.

## Abstract

Selection of regions of interest (ROIs) is often a crucial step in spatial molecular profiling and many pathology tasks, with substantial implications for research reproducibility and biological interpretability. To provide a reproducible and adaptive framework for AI-guided ROI selection, we developed a modular generalist-specialist solution across spatial profiling platforms. In a cohort comprising 55 tumor types from 160 tissue donors profiled using NanoString Digital Spatial Profiling and multiplex immunofluorescence, we first established a protein-profiling reference atlas capturing compartment-specific immune, checkpoint, stromal, and proliferation patterns. We then developed an AI Specialist Task-Oriented Model for ROI Selection (ASTROS) and tested comprehensive benchmarks considering specialist-only (ASTROS), generalist-only (PLIP/GFM), and hybrid generalist-specialist strategies, showing that the latter provides a balanced tradeoff across slide-level signal preservation, pathologist-reference concordance, within-slide placement consistency, and large-slide computational efficiency. We further demonstrated the feasibility of virtual staining for ROI preview and modular ROI placement for other spatial omics technologies, Visium and Visium HD workflows. Together, these results support our proposed framework to enable ROI selection responding to unmet needs for reducing inter-rater variability, reproducibility, and versatility in spatial profiling experiments.

## INTRODUCTION

Tumor tissue sections are essential snapshots of neoplasm evolution whose intricacies underpin cancer biology and clinical outcome^1^. Yet across pathology, genomics, and spatial omics, the biology of a whole tissue section is rarely accessible; surgical resections can span several centimeters (up to 3.7 × 2.4 cm^2^ per FFPE cassette), whereas spatial profiling platforms capture only small fractions of that area per experiment in the order of 660 × 785 μm² for NanoString GeoMx Digital Spatial Profiling (DSP)^2^, 800 × 800 µm² for multiplexed ion beam imaging (MIBI)^4^, and up to 6.5 × 6.5 mm² or 11 × 11 mm² area for 10x Genomics Visium and Xenium platforms^3^, respectively. Constrained capture areas, assay cost, and limited throughput therefore force selecting one or a few representative regions of interest (ROIs) from large heterogeneous tissue. This step underpins downstream analyses including tumor microenvironment (TME) characterization^5^, detection of immune infiltration, and quantification of immune checkpoint status^6^, cancer grading^7^, biomarker validation^8,9^, and diagnostic assessments^10^, and at the functional level can yield new insights into immunotherapy responses^11,12^, therefore, the choice of ROI directly affects study outcomes, reproducibility, and scientific validity. Yet ROI selection has traditionally been manual and expert dependent, leaving it prone to interrater variability^13^, susceptible to biases in how the whole tissue is represented^14–17^, and poorly scalable amid a growing pathologist workforce shortage^18^. Reproducible, representative ROI selection is therefore a central bottleneck for large-scale spatial profiling compounded by the fact that capture areas, assays and the biological questions they serve may differ across platforms, so no single, fixed selection rule transfers cleanly between ROI selection strategies.

Automated approaches promise to make ROI selection more reproducible by streamlining complex workflows and enabling large-scale, multi-center studies. Several studies have accordingly examined how tissue sampling represents the tumor heterogeneity. For example, Sun *et al.*^5^ derived a minimum area required to represent whole-slide images (WSIs) from the rank correlation between ROIs and WSIs, in line with established sampling principles for whole-slide tissues^16,17,20^; and, more recently, Bost *et al.*^21^ developed a systematic framework for determining the number and size of ROIs needed to accurately identify all cell phenotypes *in situ* in multiplexed images. In parallel, the accelerated progress of artificial intelligence (AI) in computational pathology has motivated classification and detection-based models for ROI selection^22–24^, including its formulation as a segmentation task through boundary update and coarse-to-fine super-pixel refinement^25^. Accounting for representativeness, Shen *et al.*^26^ localized representative patches using a pathology deformable conditional random field with a convolutional neural network backbone, Rezk *et al.*^27^ generated discriminative samples that recapitulate the population distribution via formal concept analysis, and Hossain *et al.*^28^ implemented a vision transformer for ROI detection in HER2 scoring of breast cancer. Despite these significant efforts, two major issues remain. Firstly, a systematic criterion has yet to be widely adopted with a clear goal, e.g., reducing inter-rater variability, and secondly, most of these specialist models are built for a single predefined task and cannot be readily adapted to new requirements, tissues, or platforms.

Recently popularized, generalist foundational models (GFMs) trained on a diverse corpus leverage broad contextual understanding and transfer learning promise to be adaptive across tasks^29–31^. At the other end of the generalist-specialist continuum, specialist task-oriented models (STMs) widely employed in healthcare currently achieve strong performance on narrowly defined problems^32,^ but with limited generalizability. STMs are typically fine-tuned for precise pathology tasks or disease types such as Gleason grading in prostate cancer^33^. Thus far, GFMs and STMs have largely been built and deployed as standalone pipelines rather than interoperable modules which would have their complementary strengths^34^. Therefore, harnessing both approaches synergistically is a promising strategy to characterize and represent the nuances of the TME more reproducibly; yet no modular framework has combined them such that the same generalist and specialist components can be reconfigured for different objectives and platforms, precisely what reproducible, scalable ROI selection demands.

In this work, we close this gap with a modular, generalist-specialist framework for ROI selection in histology images (Fig.1), coupling a GFM for feature extraction and representativeness^29,35^ with ASTROS, as AI Specialist Task-oriented model for ROI selection. Modularity is the organizing principle of our study design: we systematically test which components drive selection, generalist alone, specialist alone, or their hybrid integration. We demonstrated its use on clinical routine samples stained with hematoxylin and eosin (H&E) and processed with multiplex immunofluorescence (mIF) markers from a rich institutional repository of 1,116 immune-guided ROIs paired with mIF images from 160 tumor donors (GeoMX DSP, DSP cohort) spanning 55 cancer types, assembled as part of MD Anderson’s Patient Mosaic^TM^ project. We first show that AI-guided ROIs recover the immune features capture by expert-selected ROIs and by whole slides. Exploiting the framework’s modularity, we then transfer learning from mIF to H&E, generating a virtual support framework that previews AI-guided ROI selection on H&E derived virtual mIF as feasibility extension. Finally, leveraging our framework’s modularity, we reconfigure the same components through GFM, STM, and human-AI collaborations to morphology-based ROI placement in an independent cohort of 70 cases profiled with 10x Visium and Visium HD across four tumor types (Visium cohort: glioblastoma, cholangiocarcinoma, lung adenocarcinoma, and upper-tract urothelial carcinoma). Together, this framework established an integrated and flexible generalist-specialist modular strategy for representative, cross-platform, and reproducible ROI selection in spatial tissue profiling.

**Fig. 1.**
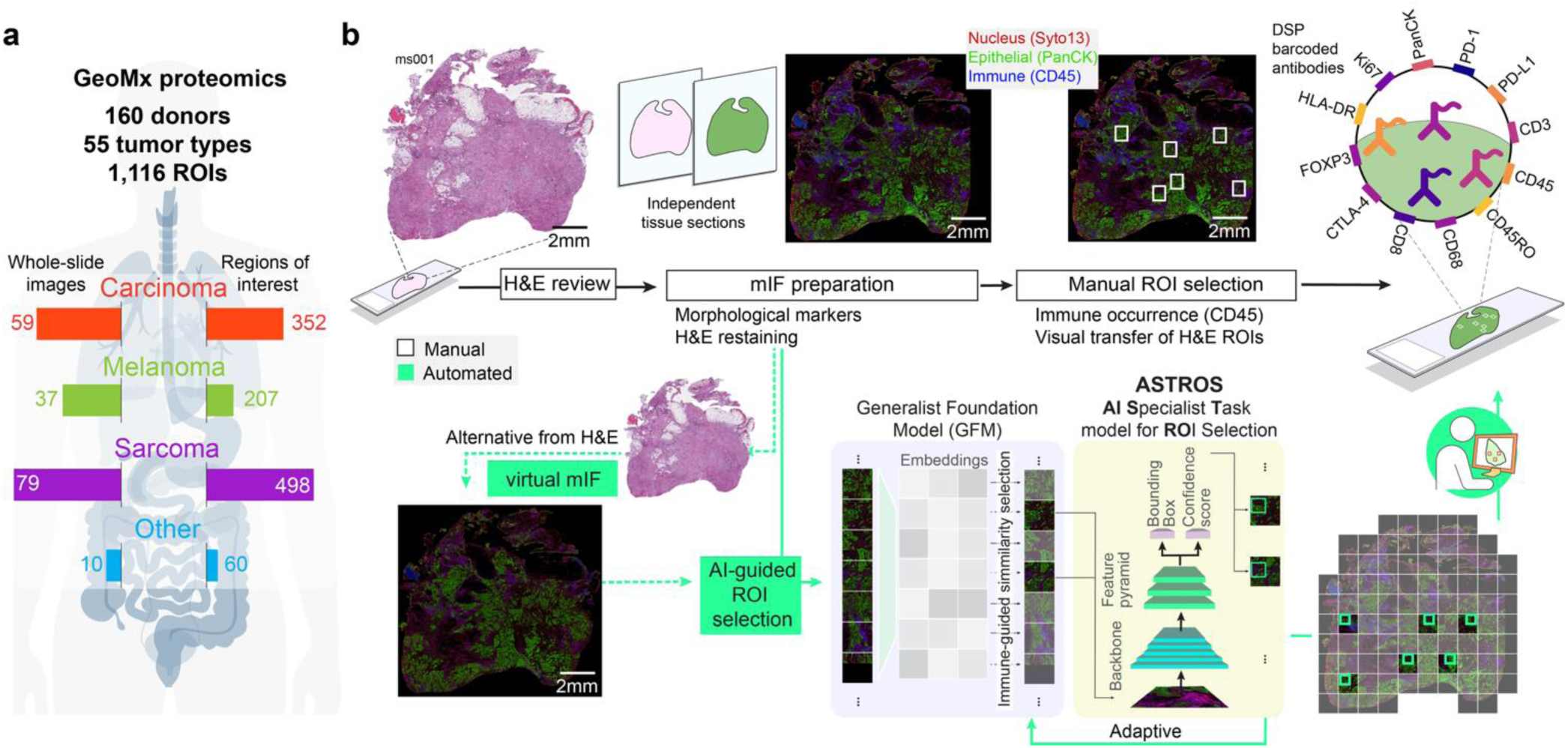
Automating the process of selecting regions for Digital Spatial Profiling. **a** We leverage a dataset from 160 donors, with tumors aggregated into categories such as carcinomas, melanomas, sarcomas, and others across different organs. **b** Schematic of the ROI selection process for spatial proteomics (NanoString’s GeoMx Digital Spatial Profiling - DSP) begins with an H&E-stained tissue section, followed by a multiplex immunofluorescence (mIF) assay from a sequential tissue cut, which is then digitized and manually annotated with ROIs. The schematic illustrates the proposed hybrid generalist-specialist approach for ROI detection for mIFs and an alternative pre-run virtual support system, built upon synthetic mIF generated from H&E re-stained tissue sections, enabling pre-assessment of the AI-guided ROI detection.

## RESULTS

### Manually selected ROIs enable comparative immunoproteomics analysis in tumors

We had retrospective access to the MD Anderson Patient Mosaic repository of 1,116 manually selected ROIs for GeoMx DSP (DSP cohort), which was utilized in this study grouped in four major categories: carcinoma, melanoma, sarcoma, and others (**Supplementary Table 1**). Manual ROIs were selected based on representativity of the whole tissue section and leukocyte occurrence identified on H&Es^36^ and CD45^+^ cells on a paired mIF tissue section. Selected ROIs in the H&E were manually transferred to the mIF image obtained from the tissue section from same block than the H&E. The mIF ROI was segmented into TME (PanCK^-^ or S100B^-^) and tumor (PanCK^+^ or S100B^+^) compartments, allowing differential expression per ROI in these two compartments (**Fig. 2a-c**), although the segmentation strategy was adjusted according to tumor types (see Methods). As a result, each ROI had an mIF image (**Supplementary Table 2**) and, after molecular profiling, barcoded DSP proteomic counts (**Supplementary Table 3**). The sequence-based proteomic panel was designed to identify targets that H&E-only morphological studies cannot capture, such as the functional status of immune cells, defining an immunoproteomic landscape across tumor types. The proteomic analysis indicated, on average, a higher proportion of TME and tumor ROIs in carcinomas and melanomas with stronger immune activity compared with sarcomas (**Fig. 2d-e**), consistent with the characterization of melanomas typically harboring dense CD8^+^ TILs and high neoantigen loads, whereas sarcoma microenvironments tend to be cold^37–39^.

**Fig. 2.**
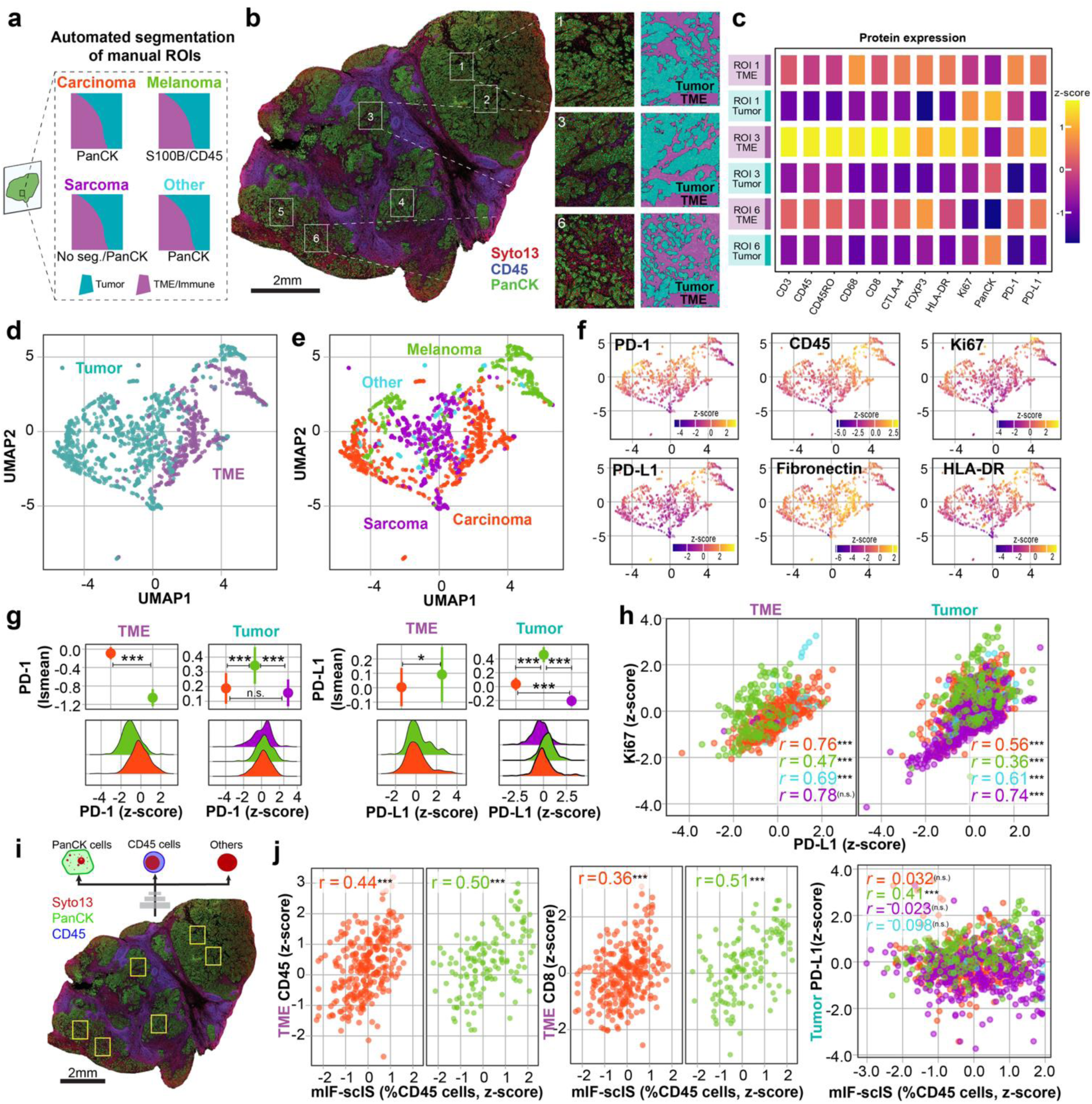
Manual ROI selection enables comparative spatial immuno-profiling. **a** Automated segmentation strategies tailored to tumor types. Proteins with immunofluorescent probes used for segmentation are indicated below each schematic, where positive expression indicates tumor, and negative expression is used to segment TME or immune regions in the case of melanomas. Only a few ROIs in sarcoma have TME segmentation (N =6). **b** Example of manually selected ROIs and intratumor heterogeneity enabling (**c**) the detection of intratumor heterogeneity of the expression of immunomodulatory proteins across the tissue in tumor (PanCK^+^) and tumor microenvironment (TME, PanCK^-^CD45^+^) compartments. The mIF-based tumor and TME segmentation masks are automatically produced during the NanoString’s DSP assay workflow. (**d-h**) Immunoproteomic landscape across (**d**) tissue compartments and (**e**) tumor types. **f** Normalized protein expression across ROIs for major proteins of interest involved in immune regulation in situ. UMAP: Uniform Manifold Approximation and Projection. **g** (top) Comparison of immune checkpoints (PD-1, PD-L1) expression (lsmean, least-square means) between carcinoma, melanoma, and sarcoma across tissue compartments. Tukey HSD p-values. (bottom) Distribution at the ROI level of PD-1 and PD-L1 expression. **h** Spearman’s correlation between Ki-67 and PD-L1 protein expression for carcinomas (λ), melanomas (λ), sarcomas (λ), and others (λ). **i** Automated cell detection based on immunofluorescence for nuclear (Syto13), epithelial (PanCK), and immune cells (CD45). **j** Spearman’s correlation between mIF single-cell score (mIF-scIS, z-score %CD45 cells) with immunoproteomics (z-score) in the TME and tumor compartments. Statistical significance (adjusted p values): p > 0.05 (n.s.), *p < 0.05, **p < 0.01, ***p < 0.001.

We observed high variability in immune modulation across tumor and TME in these tumors, based on protein expression of immune-lineage transmembrane protein (CD45), antigen presentation (HLA-DR), immune checkpoint expression (PD-1 and PD-L1), and proliferation (Ki-67, **Fig. 2f**). Across markers (**Supplementary Table 4**), inter-sample variability given by tumor type was generally substantial, especially in the Tumor compartment, where inter-class correlation (ICC) ranged from 0.57 to 0.74; the highest was Ki-67 in the tumor compartment (ICC = 0.738). In TME, ICCs were more modest for PD-1 (ICC = 0.372) but remained moderate-high for the other markers (ICCs range of 0.60 – 0.66). At the category level (**Supplementary Table 5**), Tumor Sarcoma showed the largest heterogeneity for several markers, including PD-1 (interquartile range, IQR = 1.071), PD-L1 (IQR = 0.964), and HLA-DR (IQR = 1.031), while Tumor Melanoma showed relatively high variability for HLA-DR (IQR = 1.117). In the TME, the largest IQRs were mostly seen in the Other category, especially for CD45 (IQR = 1.903) and Fibronectin (IQR = 2.097), though those estimates are based on small sample counts. Together, these results captured the overall extent of inter-sample heterogeneity of immune signaling and compartment, and the within-tumor type distribution of that variability across cancer types.

Although data availability limited our ability to study treatment response, comparative immunoproteomics indicated that carcinomas, melanomas, and sarcomas were characterized by distinct expression profiles in both the tumor and TME compartments (**Fig. 2g**). Melanomas showed higher PD-1/PD-L1 expression compared to carcinomas (PD-1: adj. p = 7.7 × 10^-6^; PD-L1: adj. p < 0.001) and sarcomas in the tumor compartment (**Fig. 2g**). In contrast, variable differences were found in the TME, with a higher expression of PD-1 in carcinomas compared to melanoma (p = 1.6 × 10^-26^) but an opposite trend for PD-L1 (p = 0.015). These patterns suggest that immune escape in the tumor region is more pronounced in melanomas than carcinomas and sarcomas, which aligns with the higher immunotherapy response rate observed in melanoma. Interestingly, we identified a consistent positive relationship between PD-L1 expression and Ki-67 across tumor types (**Fig. 2h**, Pearson’s correlation, Tumor: Carcinoma: r = 0.56, p = 1.3 × 10^-30^, Melanoma: r = 0.36, p = 1.2 × 10^-7^, Sarcoma: r = 0.74, p = 1.3 × 10^-30^, Others: r = 0.61, p = 4.9 × 10^-7^; TME: Carcinoma: r = 0.76, p = 3.1 × 10^-66^, Melanoma: r = 0.47, p = 5.3 × 10^-9^, Sarcoma: r = 0.78, p = 0.07, Others: r = 0.69, p = 7.3 × 10^-6^), indicating a strong correlation between immune checkpoint-mediated immune escape and tumor proliferation. While these findings suggest a plausible biological interplay between proliferative activity and immune evasion, further studies are necessary to confirm the mechanisms underlying the co-expression of PD-L1 and Ki-67 in the tissue. In summary, manual delineation of ROI based on mIF immune occurrence enabled a detailed and biologically meaningful comparative immunological characterization of tumors.

Next, to build on our automated framework for ROI selection, we calculated a mIF-based single-cell immune score (mIF-scIS, %CD45 cells) and tested its correlation with granular DSP immunoproteomics (**Fig. 2i**). We found significant correlations for CD45 protein present in both modalities (Carcinoma: r = 0.44, p = 2.1 × 10^-14^, Melanoma: r = 0.5, p = 5.06 × 10^-10^, **Fig. 2j**), validating the mIF-scIS estimates. Additionally, the mIF-scIS was also significantly correlated with CD8 in the TME for both carcinoma and melanoma (Carcinoma: r = 0.36, p = 1.4 × 10^-9^, Melanoma: r = 0.51, p = 1.3 × 10^-10^, **Fig. 2j**), likely related to the composition of the detected CD45^+^ cells in the mIF images.

Besides validation, the mIF-scIS score was correlated with DSP-measured PD-L1 expression in the tumor compartment only for melanoma (r = 0.41, p = 1.3 × 10^-9^), suggesting that the expression of immune checkpoint ligands in the tumor compartment could be associated with local CD45^+^ immune occurrence.

To evaluate the representativity of the WSI by manual ROIs, we compared the immune score (mIf-aIS) between these two scales. The mIF-scIS of manual ROIs and WSI are significantly correlated (Pearson’s r = 0.753, 95%CI = 0.681 – 0.810, p = 9.043 × 10^-34^). A Bland–Altman analysis showed little evidence of a fixed mean bias between methods (bias = 0.97, 95%CI = −0.33 – 2.27; paired t-test p = 0.141) with a good interclass correlation (ICC = 0.75, p = 1.54 × 10^-33^) a no effect of cancer types on the ROI and WSI scoring difference (linear regression, F_[3,174]_ = 2.05, p = 0.1082). However, the 95% limits of agreement were wide (-16.25 – 18.19), indicating substantial variability in individual-level differences between manual ROI and WSI-level mIF-scIS. These results validated the mIF automated immune scoring in manually selected ROIs and the WSI representation by ROIs, contextualizing the ROI selection for spatial immunoprofiling, providing the foundation for AI developments based on these datasets.

### Automating ROI identification based on the coexistence of immune and cancer cells

Building on insights from manual ROIs in the DSP cohort, we then sought to develop a biology-oriented automated workflow using a hybrid generalist-specialist framework and enable reproducible, large-scale analysis across diverse tumor types. To automate the process of ROI selection for DSP, we utilized the manual ROIs for this assay retrospectively accessed and devised a three-stage (**Fig. 3a**), coarse-to-fine workflow that processes mIF whole-slide images (WSIs) with a GFM for grid selection (**Fig. 3b**), and culminating in ROI selection by ASTROS, our AI Specialist Task-oriented model for ROI Selection. We designed an extensive comparison of approaches (**Fig. 3c**) to assess the advantages of using a synergistic modular generalist-specialist approach rather than its individual components standalone.

**Fig. 3.**
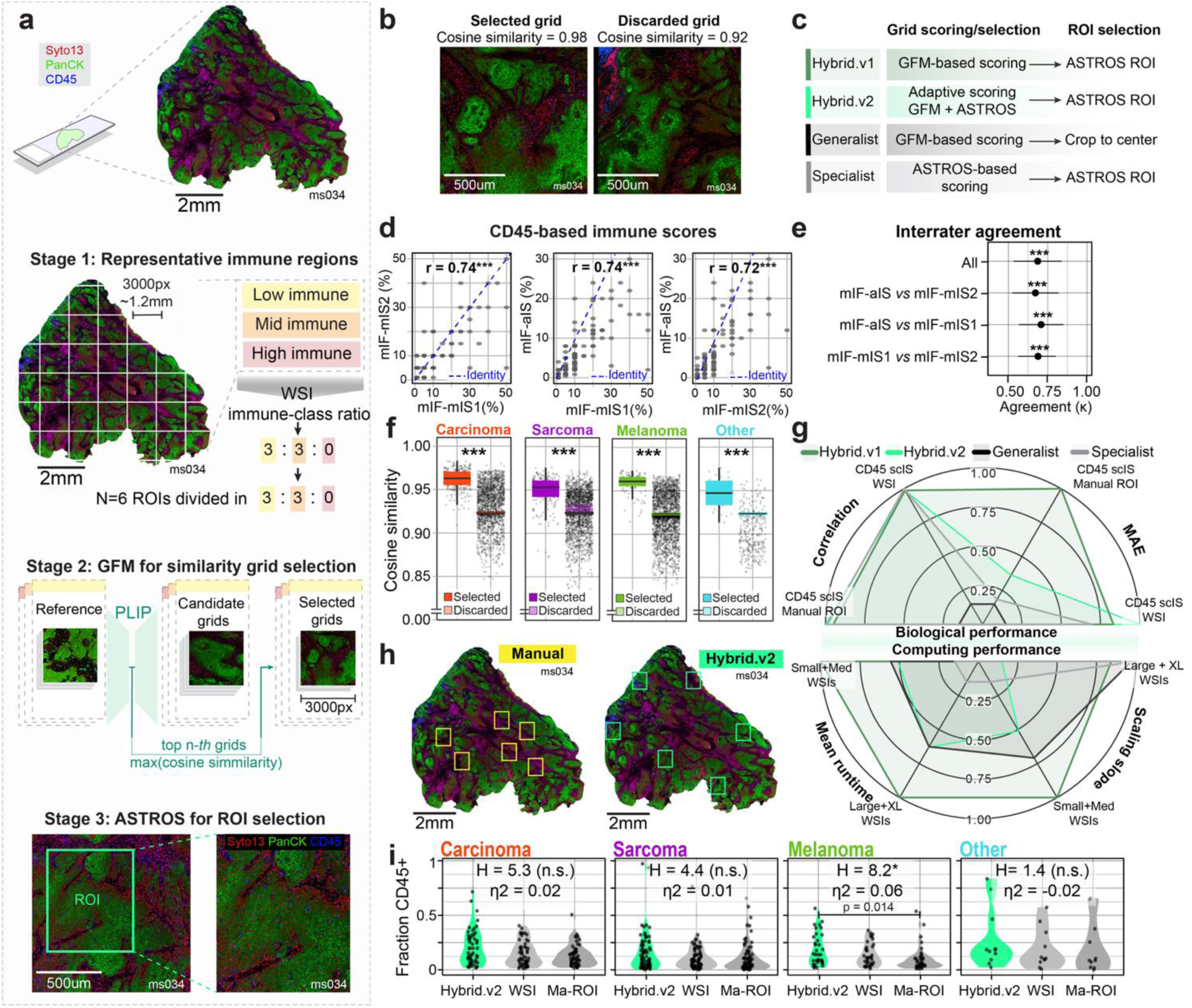
AI-guided ROI selection for multiplex immunofluorescence. **a** Workflow overview for AI-guided ROI selection with a hybrid generalist-specialist approach, including a generalist foundation model (GFM) and ASTROS, our AI Specialist Task-oriented model for ROI Selection. **b** Examples of grids selected and discarded for containing ROIs based on cosine similarity with the reference set. **c** Schema showing the four experimental AI arms considering arrangements of GFM and ASTROS for grid and ROI selection. **d** Pearson’s correlation of automated (mIF-aIS) and manual immune scores from two independent pathologists (mIF-mIS1 and mIF-mIS2). Each point represents one grid scored by all three raters. **e** Interrater agreement of categorical immune scores, Cohen’s k for orthogonal pairwise comparisons, and Light’s k for measuring the agreement across all three raters. Cut-off on continuous immune scores: 10% and 50%. Points represent k ± 95%CI. **f** Cosine similarity is based on the comparison of all possible grids and a retrospective manually selected ROIs. Boxplots show the median cosine similarity, interquartile range, and value range for selected (stronger color) and not-selected (lighter color) grids. **g** Radial plots displaying the scaled performance comparison across the four AI arms along eight axes of biological and computational performance. MAE: mean absolute error. Hybrid models tend to have higher biological performance (top half-radial plot) while Hybrid.v2 and the ASTROS model have better computation performance, with the former excelling at large and extra-large (XL) whole-slide images (WSI). **h** Examples of manual and automated ROIs in a WSI mIF. **i** Violin plot showing distribution of median single-cell immune scores (scIS) per sample across tumor types following Hybrid.v2 and manual (Ma-ROI) ROI selection and WSI-level scores. Kruskal-Wallis’s test statistics and effect size estimation (η^2^) for each comparison. Dunn test for post-hot comparison with Bonferroni-Holm correction of p-values. Statistical significance: p > 0.05 (n.s.), *p < 0.05, **p < 0.01, ***p < 0.001.

The first stage of the hybrid generalist-specialist framework included the calculation of grid-based automated immune scores (mIF-aIS) using a fast, pixel-based estimation of CD45 expression in mIFs. These scores werecompared them with two pathologists’ independent manual scores (mIF-mIS) on the same regions, showing a significant positive correlation (Pearson’s correlation, mIF-aIS vs mIF-mIS1: r = 0.74, p = 2.5 × 10^-21^, mIF-aIS vs mIF-mIS2: r = 0.72, p = 2.6 × 10^-23^), comparable to inter-pathologists’ correlation (r = 0.74, p = 5.5 × 10^-23^) (**Fig. 3d**). For categorical immune scores, using the cut-offs defined for manual selection of ROIs (10% and 50% immune cells, see Methods), the overall interrater agreement between pathologists’ scores and the mIF-aIS remains high (Light’s κ = 0.688, p = 8 × 10^-4^). Additionally, the agreement shown by pairwise comparisons of categorical manual and automated immune scores (mIF-aIS vs mIF-mIS1: κ = 0.708, p = 9.18 × 10^-23^; mIF-aIS vs mIF-mIS2: κ = 0.686, p = 1.2 × 10^-18^) was highly concordant with inter-pathologists’ agreement (κ = 0.672, p = 4.11 × 10^-19^) (**Fig. 3e**). These results indicate that image-based immune scores are a reliable estimation of immune cells occurrence as quantified by pathologists, providing a key component for automated representative ROI selection.

The second stage established the basis of representativity, we identified which and how many grids per immune category would pass to the final stage of ROI selection. This step is key to differentiating the four-arm AI approaches (**Fig. 3c**). To obtain a representative number of grids, we calculated the proportion of grids per immune category across the WSI. Then, we divided the available number of ROIs per sample (N = 6) according to that proportion, maintaining the WSI representativity across immune categories. Next, to define which grids from a WSI would pass to stage 3, we compared them with an immune-stratified reference grid set by expanding manual ROI boundaries to the same grid size as candidate grids. To calculate morphological similarity, we used PLIP^35^, a generalist foundation model (GFM) to extract embeddings from the reference grid set and candidate grids, generating a GFM score (S_GFM_) for each candidate grid as the maximum cosine similarity between its embeddings and the reference embedding space of the corresponding immune class. To verify the selection criteria, we compared the cosine similarity between selected and discarded grids, finding a higher score for the selected ones (Wilcoxon rank-sum test, W = 8.07 × 10^6^, p = 1.3 × 10^-303^, **Fig. 3f**).

This stage delivered grids that effectively captured histological patterns represented in the reference and that are ready for ROI selection (*e.g.*, **Fig. 3b**). To further refine grid selection, in our main Hybrid.v2 approach, we concurrently applied ASTROS to every candidate grid, yielding a specialist-based score S_AST_ defined as the maximum detection confidence returned for that grid. The two scores were dynamically integrated, assigning weights to S_GFM_ and S_AST_, determined based on the relative discriminative spread of each score distribution across the WSI. This formulation allowed the pipeline to adapt to slides using an adaptive synergy framework (Hybrid.v2) in contrast to Hybrid.v1, where grid selection was informed only of GFM-based cosine similarity.

In the third stage of the hybrid approaches, we developed and applied ASTROS on the selected grids. ASTROS is our AI Specialist Task-oriented model for the ROI Selection model trained with manual annotations to recognize subtle immune occurrence patterns. ASTROS was trained on 606 manually annotated ROIs across 57 cases, with a fine-tuned model incorporating an additional 222 ROIs from two independent pathologists to account for inter-pathologist variability and validated on 128 WSIs comprising 768 ROIs across carcinoma, sarcoma, melanoma, and other tumor types. In the updated pipeline, ASTROS generates final ROIs of 1,649 × 1,961 pixels^2^, matching the average manual DSP ROI dimensions.

We tested the sensitivity of Hybrid.v2 to sample sizewithin each sample and across the cohort. The within-slide standard deviation of immune estimate (mIF-scIS) decays with a power law distribution as the sample size approaches the selected size (n = 6 ROIs per sample; **Supplementary Fig. 2a**) with a similar behavior observed in the mean absolute error (MAE) (**Supplementary Fig. 2b**). At the cohort level (**Supplementary Fig. 2c-f**), we performed a bootstrap subsampling analysis (5,000 iterations per sample size) over the full pool of 1,036 Hybrid.v2 ROIs, evaluating six performance metrics: Pearson’s r, Spearman ρ, Lin’s concordance correlation coefficient (CCC), MAE, RMSE, and systematic bias against both manual pathologist and WSI grid references. At this scale, the results showed that although point-estimates of performance seem to be invariant to sampling, the confidence of their estimation significantly decays with increasing sample sizes. In summary, point estimations of immune occurrence remains invariable with sample size using our hybrid.v2 approach, but the decay decays in uncertainty with sample size indicates a robust management of variability by our framework.

We tested ASTROS’ performance with manual ROIs, reaching a mean precision of 0.969 (95%CI = 0.967 – 0.971), mean recall of 0.989 (95%CI = 0.989 – 0.991), and mAP0.5–0.95 of 0.907 (95%CI = 0.903 – 0.911) (**Fig. 3f, Supplementary Table 4**). Furthermore, to compare the performance of Hybrid.v2 with the other three AI-arms (**Fig. 3c**) and reference arms, we assessed a set of biological and computational performance metrics. The four-arm ablation isolates the contribution of each scoring component (**Fig. 3c**). Moreover, this approach demonstrated that the integrated Hybrid.v2 framework avoids the characteristic failure modes of either standalone component while preserving the ability to shift the selection objective without redesigning the pipeline. Therefore, the value of the Hybrid.v2 selection strategy lies in providing the strongest task-level flexibility across the full range of evaluation criteria simultaneously.

To estimate biological performance, we measured the MAE and Pearson’s correlation between the AI-arms and the reference arms (**Fig. 3g, Supplementary Fig. 3**) relative to immune occurrence in manual ROIs. The WSI grid census yielded a mean slide-level mIF-aIS of 16.18% (median = 13.63%, SD = 11.11%), and the manual pathologist reference 13.00% (median = 9.79%, SD = 12.26%).

Correlation with the WSI grid census was high and essentially equivalent across the three learned arms (Hybrid.v2: r = 0.857; ASTROS-only: r = 0.858; Hybrid.v1: r = 0.857; all p < 10^−51^), with GFM-only lower at r = 0.763 (p < 3.19 × 10^−35^). In terms of average immune scores relative to the WSI, the Hybrid.v1 arm was the best performing (MAE = 5.65%), followed by the Hybrid.v2 (MAE = 6.09%), ASTROS-only (MAE = 6.37%), and GFM-only (MAE = 6.89%). Additionally, the Hybrid.v2 arm achieved the highest correlation with pathologist annotation among all four arms (r = 0.624, p < 2.92 × 10^−20^), compared with ASTROS-only r = 0.622, Hybrid.v1 r = 0.6172, and GFM-only r = 0.516. This ranking held at every ROI count from n = 1 to n = 6, with the Hybrid.v1 arm consistently returning the lowest MAE against the manual reference: 8.851%, Hybrid.v2 MAE = 10.243, ASTROS-only MAE = 10.633, and GFM-only MAE = 10.708. Within-slide consistency, quantified as the coefficient of variation of pairwise intersection-over-union (IoU) across the six selected ROIs per slide, was lowest for the Hybrid.v2 arm (IoU CV = 1.424), indicating the most reproducible and internally coherent placement pattern, compared with ASTROS-only (IoU CV = 1.437), Hybrid.v1 (IoU CV =1.485), and GFM-only (IoU CV = 1.676). These results supported the ability of hybrid approaches to represent the biology and pathology of immune infiltration identified by manual ROIs and closely aligned to the WSI.

Regarding computation performance, we measured end-to-end runtime and the runtime cost of a larger dataset, estimated by the slope of runtime as a function of number of grids in a WSI (**Fig. 3g, Supplementary Fig. 4**). Mean runtimes and per-tile scaling slopes further characterize the efficiency of each arm across slide size groups. On Small+Medium slides, mean runtimes were ASTROS = 443.6 s, Hybrid.v2 = 480.8 s, GFM-only = 485.6 s, and Hybrid.v1 = 531.5 s, with scaling slopes of 3.67, 3.36, 4.09, and 3.83 s/tile, respectively. On Large+XL slides, mean runtimes were ASTROS = 1342 s, GFM-only = 1533 s, Hybrid.v2 = 1535 s, and Hybrid.v1 = 1683 s. Notably, the Hybrid.v2 arm’s per-tile scaling slope on Large+XL slides (0.64 s/tile) was less than half that of ASTROS-only (1.24 s/tile), GFM-only (1.22 s/tile), and Hybrid.v1 (1.13 s/tile), consistent with the composite scoring enabling more effective tile pruning at high tile counts. GFM-only, despite skipping ASTROS inference, achieved no meaningful runtime saving over Hybrid.v2 on large slides (1533 vs 1535 s), because unfiltered PLIP embedding on all tiles cancelled the YOLO saving. These results highlight the advantages of Hybrid.v2 for working with bigger WSI, high immune representation with a lower computational footprint, overtaking the performance when including biology and computation-relevant assessments.

After ROI prediction by Hybrid.v2, we observed cases where manual and Hybrid.v2-derived ROIs had a modest spatial overlap (**Fig. 3g**), which is likely explained by differences in the grid selection process, with the grids for manual selection being unavailable to us. Despite these differences, our quantitative evaluation indicated that the automated selection enabled by Hybrid.v2 can accurately represent the immune infiltration patterns observed in manual ROIs and WSIs despite limited overlap across most of the tumor types (Kruskal-Wallis’s test (effect size): Carcinoma: H_[2]_ = 5.3, p = 0.07, η^2^ = 0.02; Melanoma: H_[2]_ = 8.18, p = 0.017, η^2^ = 0.06; Sarcoma: H_[2]_ = 4.41, p = 0.11, η^2^ = 0.01; Others: H_[2]_ = 1.41, p = 0.495, η^2^ = −0.02; **Fig. 3h**). For melanomas, the only significant case (p = 0.017), Hybrid.v2-based ROIs’ immune scores differed from manual ROIs but not from WSIs, maintaining the consistency of the immune patterns represented by the AI-based selection. In summary, our workflow for automated ROIs in mIF images, coupling a GFM and ASTROS through a Hybrid.v2 scoring framework, resulted in Hybrid.v2 doing a precise representation of the samples, recapitulating immune infiltration patterns observed in the manually selected ROIs and WSIs, and completely reproducible.

### Developing a virtual support framework to preview ROI selection from H&Es

Given the routine availability and lower costs of H&E sections compared with mIF, we applied state-of-the-art generative AI to synthesize mIF from H&E images and run our hybrid ROI selection framework. This step enables pre-assay visualization of ROI selection using only H&E and accelerates the implementation of our hybrid generalist-specialist framework.

To create paired images for training a H&E-to-mIF image synthesis model, the mIF tissue sections used for the DSP cohort were restained with H&E and scanned. Next, these new H&Es were spatially aligned to their paired mIF images using wsireg as the image co-registration method^40^. We evaluated the performance of this co-registration method to achieve cellular-level alignment precision using a label-free quantitative approach through nuclear segmentation (**Fig. 4a**). We calculated two complementary scores on the paired and aligned binary nuclear masks: multi-scale structural similarity (MS-SSIM) and Dice. MS-SSIM was used to compare a broad set of features in the nuclear masks (luminance, contrast, and structure), and Dice scores directly measured the overlap between segmented nuclei. We found moderate to good overlap between the nuclear masks of the two image modalities (**Fig. 4b**, average MS-SSIM 0.99, median Dice scores for Carcinoma = 0.700, melanoma = 0.61, Sarcoma = 0.474, Other = 0.507), resulting in H&E/mIF image pairs readily available for image synthesis.

**Fig. 4.**
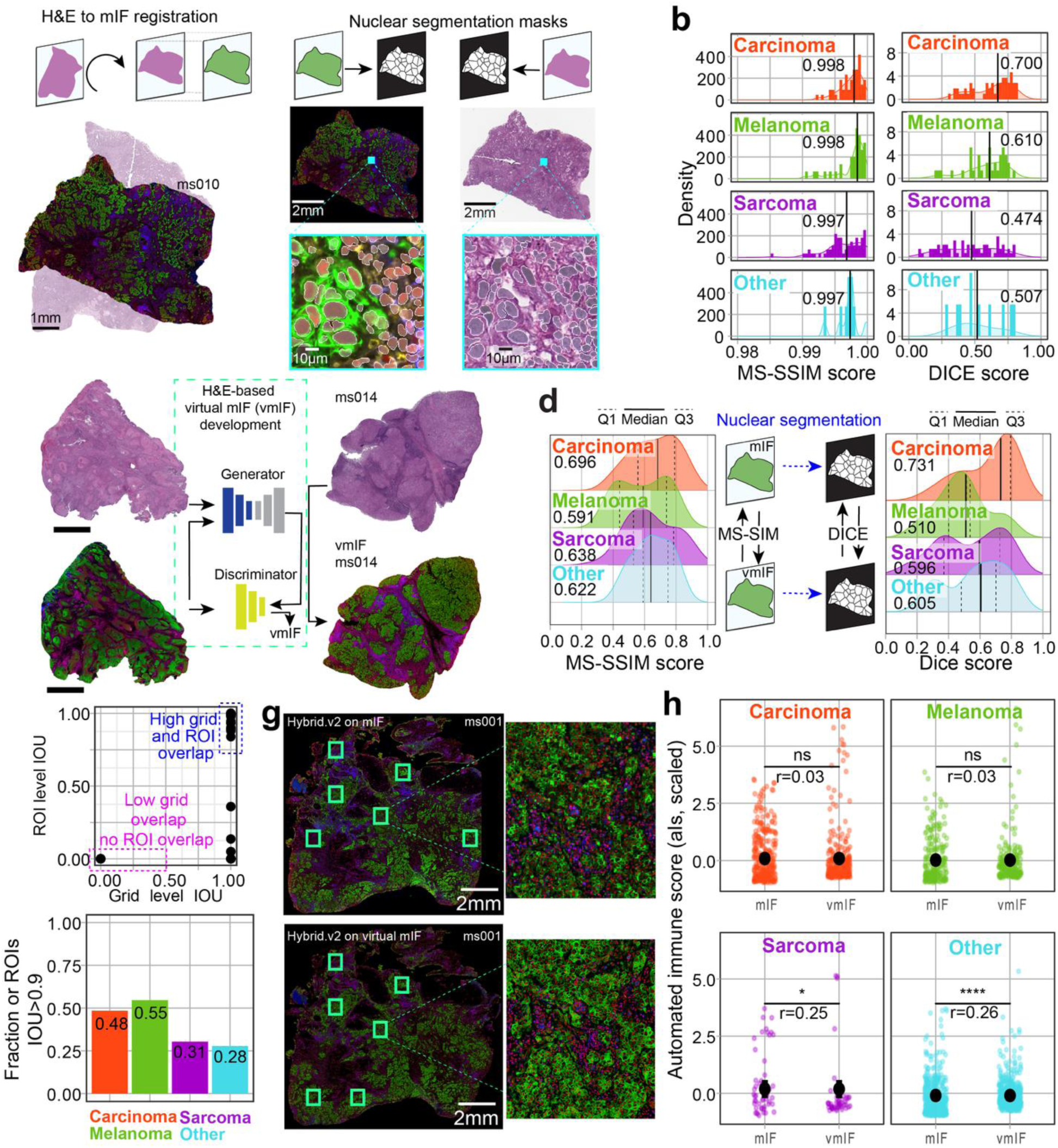
Generating a virtual support workflow for ROI selection from routine H&Es. **a** H&E and mIF images registration and approach to performance evaluation using nuclear masks. **b** Registration performance was assessed by generating nuclear masks in case-matching mIF and H&E images and evaluating their multiscale structural similarity (MS-SSIM) and Dice scores. Vertical segmented lines and adjacent values indicate the median score value. **c** Synthesis of virtual mIF (vmIF) using Pix2Pix and paired aligned mIF and H&E images. **d** Performance evaluation of vmIF synthesis in test dataset, MS-SSIM score calculated on the images, and Dice scores calculated for nuclear masks. Median score values are shown in each plot. (**e**) Overlap between grids and automated ROIs in mIF and vmIF, below IOU at grid level, no overlap is achieved between ROIs, above IOU = 0.5, the overlap is the highest (IOU-ROI=1.00), reflected in (**f**) nearly 50% of ROI are correctly predicted in the vmIF for carcinomas and melanomas, and slightly lower for the rest of the samples. **g** Example of AI-guided ROI selection on mIF (top) and vmIF (bottom) with a zoom-in to one of the matching ROIs. **h** Quantitative comparison (Wilcoxon test) at the slide level of the average single-cell immune score (scIS) between ROIs selected in mIF and vmIF. Range point indicates median values, and vertical lines show an estimated pooled range. Power analysis was estimated with rank-biserial correlation (r). Statistical significance: * < 0.05, **** p < 0.0001, p > 0.05 (n.s.).

Using the mIF/H&E aligned pairs, we trained Pix2Pix^41^, a conditional generative adversarial network enabling virtual mIFs (vmIF) synthesis from H&E. To evaluate the performance of the synthesis, we calculated MS-SSIM scores for the virtual mIF (vmIF) with the actual mIF (**Fig. 4c**). Additionally, Dice scores were calculated for nuclear segmentation masks in vmIF and mIF. The evaluation of the vmIF synthesis resulted in satisfactory performance with a median MS-SSIM of 0.644 (95%CI = 0.62 – 0.67) and a median Dice score of 0.60 (95%CI = 0.56 – 0.63). Subsequently, we applied the hybrid generalist-specialist approach on the vmIF as described for mIF by using the PLIP GFM for feature extraction and grid selection, and ASTROS, our specialist model for ROI selection.

We compared the automated selection of ROIs on the vmIF with its mIF counterpart in terms of spatial overlap and immune composition. Due to potential misalignments between mIF and vmIF, the grid step could have an offset. To address this, we established pairwise comparisons between each mIF-ROI and all vmIF ROIs within a slide, calculating their overlap and including the overlap in their corresponding grids. For low intersection over union scores at the grid level (IOU < 0.5), there was no matching between mIF and vmIF ROIs. However, with the grid’s IOU > 0.5, the ROI-level IOU score reaches the maximum overlap (**Fig. 4e**). This resulted in approximately 50% of mIF ROIs showing high overlap with the vmIF in carcinomas and melanomas, with a smaller fraction observed in sarcomas and other tumor types (**Fig. 4f**). Despite limited overlap in some cases (*e.g.*, **Fig. 4g**), at the slide level, the mean immune score in vmIF ROIs (vmIF-scIS) did not differ from the scores in the mIF ROIs (mIF-scIS), suggesting these were both comparable subsets. (**Fig. 4h**, Wilcoxon test, p > 0.05). We identified that, despite the positioning of vmIF ROIs, using ASTROS helps to select consistent immune patterns in both the virtual and actual mIFs, supported by a significant correlation between mIF and vmIF immune scores at the slide level (Spearman’s r = 0.48, p = 2.46 × 10^-8^). Further support was reached after statistically controlling for registration and virtual staining performance, with the correlation increased by 21.82%, reaching a Spearman’s partial correlation of 0.587 (p = 3.84 × 10^-8^). These analyses delivered certainty on end-to-end automated ROI selection only from H&E, thus serving as a functional pre-assay approach allowing low-cost generation of virtual mIF for ROI selection.

### Adaptation of the framework to spatial transcriptomics

Having shown the suitability of automated ROI selection in the DSP workflow, we adapted our hybrid generalist-specialist framework for ROI selection on 10x Genomics’ spatial transcriptomic platforms (Visium and Visium HD). We used retrospective data from H&E-stained WSIs from four tumor types, Visium cohort: glioblastoma (GBM), cholangiocarcinoma (CCA), lung adenocarcinoma (LUNG), and upper-tract urothelial carcinoma (UTUC). To showcase the flexibility of our approach, we explored three scenarios with different strategies for ROI selection, tailored to meet various needs. Finally, we deployed our framework with a pathologist’s oversight for prospective ROI selection in CCA, which was used as input for Visium HD. For these cohorts, we focused on the performance of ROI selection and the interpretability of those.

The first scenario reflected a common need to select ROIs that capture representative tumor areas while maximizing viable tissue coverage within the capture region (**Fig. 5a**). For this purpose, we implemented a GFM-based approach that encodes critical tissue features for ROI selection, such as viable tissue with low hemorrhages and necrosis. Each WSI was tiled and passed through UNI^29^, a state-of-the-art GFM, for embedding extraction. With the embeddings, we trained a classifier using pathologists’ annotated ROI as labels from LUNG, CCA, and UTUC and tested it on GBM. We fitted a logistic regression model to generate the probability of each tile being within the ROI. We found that automated ROI selection using a GFM generated accurate results for GBM tissues, visualized by an ROI probability map (**Fig. 5b**), and quantitatively supported by the high overlap with manual ROIs (**Fig. 5c**, N = 27, median Dice ± SD = 0.83 ± 0.32, median IOU ± SD = 0.71 ± 0.33). Therefore, our GFM-based method leveraged critical contextual information and provided a fast and generalizable ROI selection approach for spatial transcriptomics assays.

**Fig. 5.**
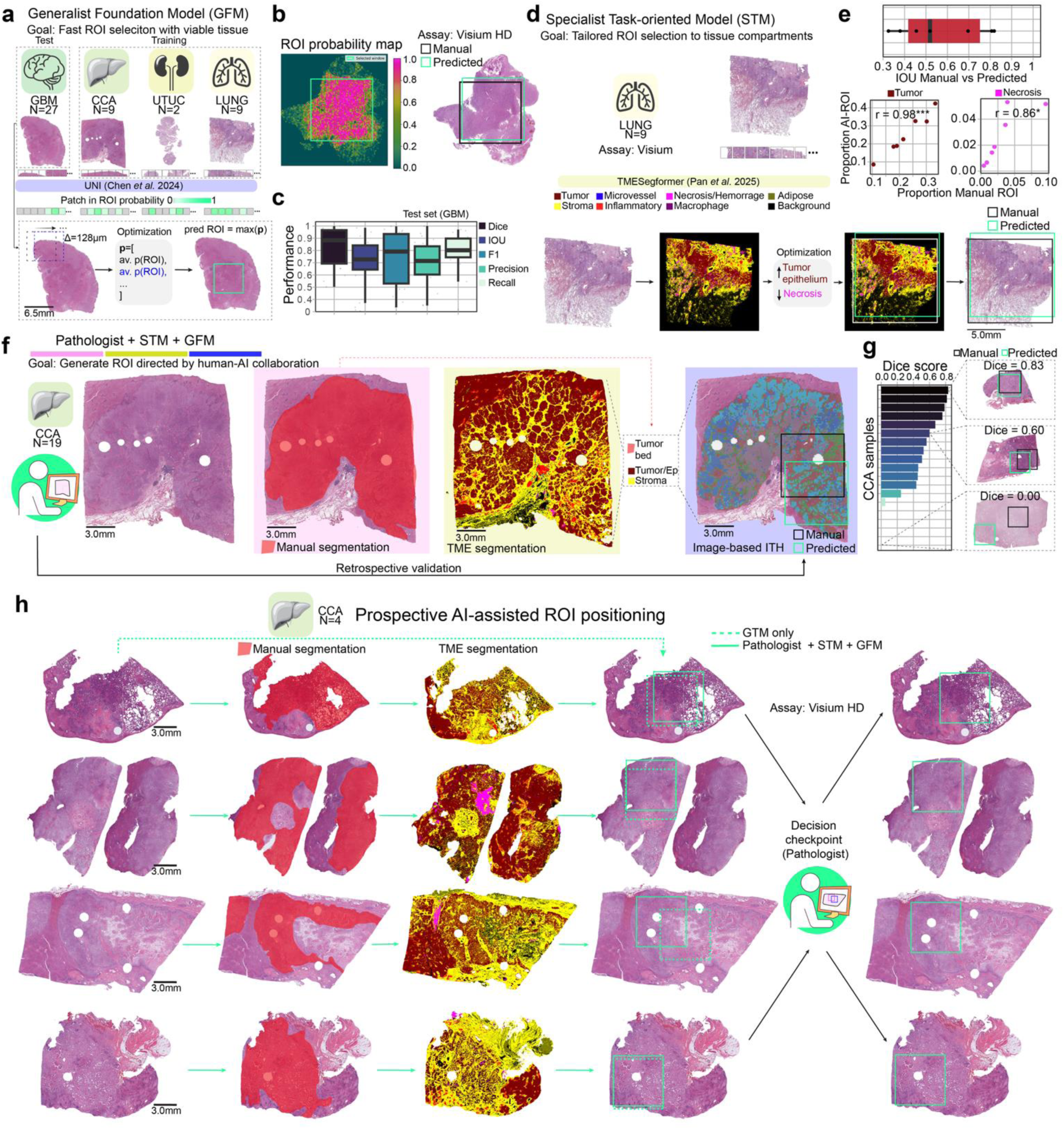
Adaptation of the generalist-specialist framework for H&E ROI selection in 10x’s Visium and Visium HD spatial transcriptomic platforms. **a** A fast implementation with a generalist foundation model (GFM) to extract embeddings from a WSI with a systematic sampling from each WSI through a sliding window, optimizing cross-tumor tissue viability. **b** ROI probability map example with the accompanying final prediction and manual annotation overlaid on an H&E. **(c)** Performance metrics for the ROI prediction in the test set glioblastoma (GBM). Boxplots indicate median performance (horizontal thick line), interquartile range (box limits), and data range (whiskers). **d** A tailored approach using TMESegformer, a specialist task-oriented model (STM) pretrained for TME segmentation, for ROI selection by maximizing the presence of tumor tissue and minimizing necrotic components. **e** The automated ROI selection is correlated (Pearson’s correlation) with the manual ROI selection’s proportion of tumor tissue and necrosis. Statistical significance: p > 0.05 (n.s.), *p < 0.05, **p < 0.01, ***p < 0.001. **f** A more complex use case involves a pathologist’s input to define a region, followed by an STM for tissue semantic segmentation, and finally a GFM that encodes tissue tiles into their embeddings, enabling a computational approach to intratumor heterogeneity. **g** Performance of retrospective application of the synergic modular approach for ROI selection and **(h)** prospective proof of concept in the position of an ROI assisting pathologist’s selection for four samples processed for Visium HD.

The second scenario addressed specific requirements for including or excluding certain tissue components, such as necrosis. With that aim, we utilized TMESegformer^42^, a pre-trained specialist task-oriented model (STM) for tumor segmentation, for ROI selection in our LUNG cases (N = 9). The model was trained using breast and lung cancer samples, and can segment tissues into ten categories, including epithelium/tumor, necrosis, and inflamed regions, among others^42^. After the generation of WSI-level segmentation mask, the region with the highest tumor/epithelial-to-necrosis ratio was selected using a moving window approach (**Fig. 5d**). We compared the results with pathologist’s manual ROIs showing a moderate-to-high overlap (median IOU ± SD = 0.45 ± 0.21, **Fig. 5e**), with a significant correlation in the tumor/epithelium (Pearson’s correlation r = 0.98, p = 6.21 × 10^-6^) and necrosis (Pearson’s r = 0.86, p = 0.003) composition. Cases with lower overlap (*e.g.,* IOU = 0.34; **Supplementary Fig. 2**) can be attributed to the stringency of the automated approach, minimizing necrosis, while manual selection may have been more permissive. The results of this scenario demonstrated that STMs, specifically TMESegformer, are more informative and explainable alternatives for automated ROI selection, offering more flexibility for specifications.

Finally, the third scenario involved a more complex study with needs and design that requires human-AI collaboration. We merged the previous two approaches into a hybrid specialist-generalist approach guided by coarse pathologists’ manual segmentation (**Fig. 5f**). This is particularly useful when only a tissue compartment is of interest, and there is no specialist model for automated segmentation, thus requiring domain expert annotation. In our implementation, a pathologist annotated the tumor bed as the macroscopic area of interest, which was then further refined by the hybrid STM-GFM method.

TMESegformer was used for tissue segmentation to distinguish tumor/epithelium and stromal compartments, followed by a GFM application to generate a morphological representation of the pathologist-annotated region and candidate ROIs obtained through a sliding window. The candidate ROI with the highest cosine similarity with the tumor/epithelium and stromal compartments in the pathologist-annotated region was selected as the ROI. In the evaluation set (CCA, N = 19), we observed variable performance compared with manual ROI selection (median Dice ± SD = 0.46 ± 0.31, **Fig. 5g**). While some cases demonstrated effective ROI selection (Dice score > 0.5), others with more homogenous tissue types remained challenging. In such cases with limited heterogeneity, multiple ROI placements were optimal, leading to discrepancies between manual and automated ROIs.

These three scenarios illustrated the advantages and limitations of single-model approaches and the benefits of a modular framework for ROI selection. Further, as proof of concept, we applied the framework to prospectively select ROIs in CCA samples (N = 4) used for Visium HD (**Fig. 5h**).

Scenarios I and III were applied to this set. A pathologist evaluated the automated ROIs and decided whether to keep one of the automated ROIs or manually annotate a new ROI. In three cases, the pathologist approved the automated ROIs predicted with scenario III (manual tumor bed annotation + STM + GFM). In only one case, both predictions were too similar, so scenario III was retained for consistency. For these four cases, the predicted automated ROIs were used as input for the assay. With this application, we demonstrated our workflow readiness, synergistically combining generalist foundation models, specialist task-oriented models, and domain-expert input for ROI selection through a transparent, reproducible, and scalable workflow that can be adapted to different hypothesis-driven scenarios.

## DISCUSSION

The automation of ROI selection for tissue assessment has been identified as one of the AI applications that will be most likely used by pathology labs by 2030^43^. Ideally, automation should enable a consistent and reproducible capture of relevant biological patterns in tissues, while maintaining whole-tissue section representativity and adaptability across diverse tissue profiling platforms and project needs.

Previous works^22–24,27,28,44^ have demonstrated how AI-based approaches can contribute to ROI selection through single-model approaches, such as convolutional neural networks or vision transformers. However, the deployment of niche computational pathology models hinders swift adaptations to heterogeneous histological data. Foundation models have emerged as a potential solution to bridge the gap^29^. New computational frameworks are needed to leverage the powerful contextual capacities of foundational models while incorporating the task-oriented performance of specialist models to facilitate reproducible scientific discoveries. In this work, we showed the feasibility of a generalist-specialist modular integration for ROI selection as an explainable and adaptive approach, expeditiously applicable to different image modalities, research questions, and spatial omics assays.

We defined representativity, flexibility, and scalability in a task-specific manner. Representativity refers to preservation of WSI and manual ROIs histological features relevant to the automated ROI selection. Flexibility accounts for the ability to combine generalist tissue-similarity filtering with specialist ROI localization in a modular fashion. Finally, scalability refers to the ability to apply this workflow across large WSIs with measurable runtime and annotation-efficiency tradeoffs. For multiplex immunofluorescence, we described a three-stage process that leverages published foundational models and a custom-designed STM, ASTROS, which emulates human selection, capturing broader patterns across samples, followed by targeted optimization and selection of specific ROIs. While ASTROS was trained with extensive manual annotations, as we demonstrated, a relatively small manual reference set may be sufficient due to the contextual information-processing capability and the growing performance of GFMs^29^. This will depend on the complexity of the ROI selection criteria.

Alternatively, to reduce reliance on extensive annotations, less supervised approaches and contrastive learning may be introduced or morphology to omics translations^45^ with the associated challenges of representing morphology and omics in a shared embedding space^46^. Although subjectivity can be introduced through manual annotations used for training, it remains essential to validate expert-domain annotations and anchor the selection on established protocols to ensure reliability. As we demonstrated in this study, manual selection of ROIs guided by community-established guidelines for immune presence in tumor tissues^36^ helps to capture immunoproteomic activation patterns. Through automation of this process, our workflow can reduce labor, particularly for double ROI selection (first in H&E and then on mIF), while accounting for the variability and complexity across tumor types. The value of a modular generalist–specialist framework lies, henceforth, in allowing these objectives to be combined, tuned, and extended without constraining the ROI selection task into a single rigid model.

We primarily focused on the analysis of the immune-based NanoString’s GeoMx DSP platform; however, our framework can be easily tailored to diverse scenarios due to its modular layout. For example, we expanded the application for ROI selection for the Visium platforms, which showcased the application to other tissue compartments as an explainable approach while keeping the expert in the loop and drawing their attention to these areas. We showcased diverse conformations of GFMs, working in conjunction with STMs, and incorporating domain-expert inputs. We showed that although the GFM-based approach was an expeditious implementation, some use cases may require more explainability and specificity in the ROI selection, as elaborated in the STM and human-AI collaboration scenarios. Keeping the experts in the loop, in our opinion, could further facilitate the adoption of this framework. The expansion to other platforms with different requirements, such as *in situ* hybridization or immunohistochemistry for clinical biomarkers, remains to be explored in future studies. For addressing these application gaps, we anticipate that an expansion can be achieved by incorporating more domain knowledge, GFMs, and STMs, integrating multiple modalities (*e.g.* ^47^). For example, additional modules could include tailored models for single-cell classification, spatial pattern detection, image retrieval, or even quality control^48^ to optimize ROI selection, completely tailored to platforms beyond DSP and Visium’s requirements.

One aspect that partially escaped our evaluation is the effect of sample size. This is particularly relevant when multiple candidate ROIs are present in a tissue. Sampling theory suggests that the precision of estimating population parameters increases asymptotically with sample size, as demonstrated in ROI selection^5,21^ and cohort-level simulations^49^.Our results point in the same direction. In recent efforts, Baker *et al.*^50^ proposed a power analysis framework that explores the design of spatial molecular assays by incorporating parametrization based on cellular distribution, with applications such as the detection of cell-cell adjacencies as a function of sampling effort. For spatial pathomics assays, it is challenging to test the effect of sample size (*i.e.*, the number of ROIs) on omics patterns due to the associated experimental costs, thus facing a critical tradeoff between the number of samples and the number of ROIs per sample. Platforms like Patient Mosaic, as the DSP cohort included here, have faced this challenge by prioritizing sample size while keeping WSI-level representativeness. For our case, the modest sample size and patient-history heterogeneity limit more robust molecular conclusions, which should be taken into consideration for gauging the molecular descriptions between tumor types. For future development, a more direct sample-size evaluation could be accommodated with a careful experimental design and prospective sample size analysis that incorporates answering biologically relevant hypotheses and tests the suitability of automated approaches. With the advent of spatial omics, public resources will enable the generation of comprehensive datasets, allowing for more robust statistical evaluations based on morphological representation across different tumor types and platforms.

Although our hybrid generalist-specialist framework performs ROI selection in relevant areas for spatial omics, some challenges remain for complex histomorphologies, such as well-differentiated liposarcomas, where their adipocytic structures limit the direct transferability of a generic tool. These challenges could be exacerbated during the synthetic immunofluorescence stage, as H&E and mIF might share limited information in the latent feature space, thus preventing accurate synthesis. For cases with complex and infrequent morphologies, a fine-tuned version could be deployed together with a tailored immunofluorescence panel and reference grid set. This approach would align with generalist approaches as a starting point and, if needed, fine-tuned or specialist approaches.

In summary, although GFM and STMs are often presented as competing views due to their different approaches to automated data analysis, we demonstrated a pioneering implementation where a robust, modular AI-Pathology ecosystem can successfully leverage their strengths and enable their synergy. Additionally, integration with domain-experts’ knowledge with this hybrid system design both advances scientific reproducibility, improves data analysis scalability, and has a profound impact on clinical utility in domains such as biomarker discovery and deployment by eliminating technical noise introduced by manual tasks. We demonstrate that synergistic approaches are adaptive and can be tailored to develop reproducible workflows for ROI selection in spatial omics, with swift applications in digital pathology, thereby enhancing readiness and facilitating the discovery of fundamental immune profiling crucial for cancer biology.

## METHODS

### Ethical statement

All human studies were conducted following the relevant ethical guidelines and approved by the appropriate committees at MD Anderson Cancer Center. Written informed consent was obtained from all participants or waived for deceased patients (Institutional Review Board (IRB) #2014-0938).

Surgically resected tumor tissue for research was collected as part of standard of care procedures, analysis performed under #PA12-0305 for spatial proteomics in tumor, and data aggregated under #2022-0404. Retrospective data access was granted and analyzed under the IRB #2023-0114 at the University of Texas MD Anderson Cancer Center. For spatial transcriptomics, data processing followed the IRBs #2016-0867/#2014-0820 (glioblastoma), #PA16-0723 (cholangiocarcinoma), #PA12-0305 (upper tract urothelial carcinoma), and #PA14-0241/Lab07-0233 (lung).

### Dataset description

This study had retrospective access to tissue scans and sequence-based immunoproteomics profiling from 160 tissue donors from studies affiliated with the Patient Mosaic^TM^ program at MD Anderson Cancer Center (DSP cohort**, Supplementary Table 1**). Each case was classified according to the diagnosis in carcinoma (55 cases, 34.4%), melanoma (28 cases, 17.5%), sarcoma (67 cases, 41.9%), or other (10 cases, 6.2%), with more complex diagnoses, such as mesothelioma, chordoma, or uterine carcinosarcoma. More granular pathological classification was available (**Supplementary Table 1**); however, the investigation of pathological subtypes was not within the scope of this study.

Tissue blocks were obtained from each patient’s tumor, sectioned for pathological quality control using Hematoxylin & Eosin (H&E) staining. Then, a serial section was prepared for multiplex immunofluorescence (mIF) with a morphological antigen panel (nuclear: Syto13, epithelium/cancer cells: PanCK, melanoma cells: S100B, and immune: CD45) (**Supplementary Table 2**). This morphological panel was validated following^2^. The mIF tissue was scanned according to the NanoString’s Digital Spatial Profiling (DSP) protocol at 40× magnification (image resolution = 0.4 mm/pixel). We obtained retrospective access to mIF whole-slide images (WSI), manually annotated ROIs on mIF, and the resulting roi × counts matrix from DSP for the immunoproteomics panel (**Supplementary Table 3**).

### Manual selection of ROIs

Quantitative evaluation of immune infiltration, adapted from the TILs Working Group^36^, guided manual ROI selection for the DSP assay in the DSP cohort. Experienced pathologists (AS, LS-S) screened each H&E WSI and calculated an immune infiltration score (percentage of lymphocytes) across a regular grid (3000 × 3000 pixels^2^). The scores were grouped into three immune categories: low immune score (< 10% lymphocytes), mid immune score (10-50% lymphocytes), and high immune score (> 50% lymphocytes). To target a selection representative of the whole-slide tumor immune infiltration, a ratio of immune categories was calculated and maintained for allocating the total number of possible ROI according to this ratio. Then, the pathologists visually transferred the selection from H&E to mIF, selected grids with viable tumor and immune infiltration in the corresponding classes. Within each chosen grid, the final ROI was drawn to represent the grid’s characteristics and exclude areas with low cellularity or non-target regions that produce expression artifacts (*e.g.,* hemorrhages). On average, six ROIs per sample were manually annotated, with regular or irregular shapes, as representative regions of the whole-slide immune infiltration. Only rectangular ROIs were retained, totaling 1,116 ROIs (median size = 512 × 512 mm^2^) across 169 WSIs. The manual selection was completed before computational development. At the time of this study, the information regarding annotations and ROI preselection on H&E and grids on mIF leading to manual ROIs was not available.

To assess the representativeness of manual ROI of WSI CD45 expression (single-cell immune score, scIS) we calculated scIS for each manual ROI and for a moving window (3000×3000 pixels^2^) across each WSI. The median values of each scale, ROI and WSI were taken for assessing statistical significance (Pearson’s correlation and paired t-test) and for a Bland-Altman analysis. Additionally, we calculated the interclass correlation between both scales.

### Proteomic digital spatial profiling

After manually selecting ROIs, the tissue was processed according to the manufacturer’s recommendations using the NanoString’s nCounter® system and the MD Anderson’s immunoprofiling platform protocol for sample preparation^2^. A customized barcoded immune proteomic panel was the target, in addition to the housekeeper and background proteins for quality control assessment (**Supplementary Table 3**).

Based on protein expression in mIF, each manually selected ROI was automatically segmented for tumor or TME, resulting in protein counts for each segment per ROI (**Fig. 2a**, **Supplementary Table 2**). The strategy to spatially profile these tumors consisted of obtaining quantitative biomarker information of the tumor and the surrounding TME. Due to variability in morphological features and availability of tumor markers, the ROI selection and segmentation were approached distinctively by tumor type. In carcinomas, the ROIs were placed in areas with tumor and surrounding TME, including immune cells and tumor stroma. The ROIs were segmented into tumor (panCK^+^) and TME (PanCK^-^).

For melanomas, as they usually display scant stroma, the ROI was placed in tumor areas with the presence of immune cells, and the segmentation for tumor was performed using S100B^+^ and the TME using CD45^+^. In the case of sarcomas, due to their wide heterogeneity of morphological features and biomarker expression, the ROI was placed distinctively in tumor areas and in TME (surrounding stroma) with no further segmentation strategy. This reflected the morphological arrangement in sarcomas as sheets of tumor cells and non-malignant stroma, sometimes observed surrounding tumor areas.

The protein count matrix across ROIs and samples was processed following NanoString guidelines. For normalization, two metrics were assessed for technical performance: housekeeper geometric mean and IgG geometric mean. To choose which geometric mean to consider for the calculation of the conversion factor, we evaluated the correlation within the background (Rb IgG, Ms IgG1, and Ms IgG2a) and housekeeper (S6, Histone H3, and GAPDH) protein classes, keeping the geometric mean from the group with the strongest average correlation. The background geometric mean was used for normalizing protein counts across ROIs and samples. After normalization, protein counts were log-transformed (base 2) and scaled (z-score) for principal component analyses, using 33 principal components for Uniform Manifold Approximation and Projection for Dimension Reduction (UMAP), resulting in two dimensions for visualization.

For each marker and tissue compartment, we fitted a linear mixed-effects model with category as a fixed effect and sample as a random intercept. The overall effect of category was assessed by analysis of variance of the fitted mixed model, and estimated marginal means (EMMs) were computed to obtain model-based category means. Pairwise differences between categories were evaluated using post hoc contrasts of the EMMs with multiple-testing corrections. Model-derived variance components were used to estimate the between-sample variance, residual variance, and intraclass correlation coefficient (ICC). To further characterize category-specific heterogeneity, we calculated sample-level mean values for each marker and summarized their distribution within each category using the mean, standard deviation (SD), interquartile range (IQR), and coefficient of variation (CV). Spearman’s correlations were estimated for Ki-67 vs PD-L1 expression across tumor types and tissue compartments. For comparison involving more than one contrast, Bonferroni-adjusted p-values were calculated.

### Correlation between mIF-based and sequence-based CD45

Since barcode-based immune proteomics is heterogeneous across ROIs and tumor types, we then assessed the association between mIF-based immune infiltration and barcode-based immune proteomics. To quantify immune infiltration in mIF-ROIs, we utilized an in-house semi-supervised deep-learning model^51^ that enables single-cell classification into three major classes: immune cells (CD45^+^), cancer cells (PanCK^+^/SB100B^+^), and other cells (CD45^-^ and PanCK^-^/SB100B^-^). In short, this model used a partially labelled dataset to train a detector that is supplemented with pseudo-labels within an iterative training loop. The pseudo-labels were then used to fine-tune the model’s learning. For prediction, the model takes mIF images (Syto13, PanCK/ SB100B^+^, and CD45) and returns the class for each cell detected. This model enabled us to obtain estimates of immune cell occurrence (percentage of CD45 cells, single-cell immune score scIS) that we correlated with protein counts. All variables were log2-transformed and independently scaled.

To compare the expression of DSP proteins between tumor categories, without removing inter-ROI heterogeneity by simply calculating the mean value per case, we implemented a generalized linear model with a nested structure. These types of models are the most appropriate statistical approach in cases with more than one observation per biological replicate, such as multiple ROIs per case. Once the model was fitted to the data, the least-squares means were calculated and plotted. We also investigated inter-case variability stratified by cancer type by separating models by tumor type allowing us to compare cancer types through estimated case-level variance partition coefficients, and likelihood-based confidence intervals. To evaluate statistical significance in the comparison between tumor categories, a Tukey HSD test with a 95% confidence level was used, allowing for the adjustment for multiple comparisons when more than one pairwise comparison is included. Overall, we performed Spearman’s or Pearson’s correlation as specified in each case to test the association between continuous variables. The association between categorical immune scores was evaluated with interrater agreement. The comparison of continuous variables across categorical independent variables was done with the Wilcoxon test. Post-hoc comparisons were done if needed, correcting p-values for multiple comparisons. Across all comparisons and statistical tests, we maintained α = 0.05.

We compared the median immune scores (scIS) resulting from AI-based and manual ROIs with WSI-level estimates, with a Kruskal-Wallis test between all three methods. We estimated effect size by including η^2^ values. In the case of a significant Kruskal-Walli’s test (p < 0.05), a Dunn test was used for post-hoc pairwise comparisons, with p values adjusted using the Bonferroni-Holm correction.

#### Automating ROI selection on mIF images

Our methodology has three key stages for automated ROI selection in mIF images (**Fig. 3a**). First, we pre-processed the mIF WSI to quantify immune infiltration using a CD45 expression immune score.

Next, a grid was established, and representative, immunologically stratified regions were selected based on GFM-based feature similarity between the grid and a retrospective dataset annotated by pathologists. Finally, for the automated ROI selection on the selected grids, we trained and applied ASTROS, our AI-based Specialist Task-oriented model for ROI Selection.

### Automated immune scoring

To accurately and efficiently estimate immune infiltration in WSIs, we calculated a pixel-based automated immune score (aIS) on mIF. First, we manually excluded regions with artifacts, such as those produced by hemorrhages and blood vessels that could confound the analysis. Then, for precise tissue segmentation, we employed hole-filling techniques to construct a comprehensive contour mask, enabling us to distinguish between glass and tissue regions within the tissue sample. We defined stroma as the region within the tumor devoid of PanCK^+^/SB100B^+^ cells, artifacts, or necrosis. Next, in 1.2×1.2 mm^2^ (3000×3000 pixels^2^) non-overlapping grids throughout the mIF WSI, mIF-aIS is determined for each grid by calculating the ratio of immune pixels (CD45^+^) to the total tissue pixels (CD45^+^ + Syto13^+^ + PanCK^+^/SB100B^+^ pixels). We excluded from this scoring those grids with > 50% background pixels.

To evaluate the accuracy of the mIF-aIS for quantifying immune infiltration, we compared it with manual immune scoring (mIS) performed by two independent pathologists blinded to the aIS. The mIF-mIS score was estimated as the percentage of CD45^+^ cells over the total number of cells. We evaluated the association between mIF-aIS and mIF-mIS with Spearman’s correlation for continuous values. For orthogonal pairwise interrater agreement (two pathologists’ scores and aIS) on categorical scores, we calculated Cohen’s k (R package: psych). Categorical cut-offs were used as previously described: low immune occurrence score < 10%, mid immune occurrence score > 10% and < 50%, and high immune occurrence score > 50%. We calculated the overall interrater agreement using Light’s k (R package: psych).

### Grid selection representative of the WSI cancer-immune coexistence patterns

After calculating mIF-aIS for each grid in a WSI, we classified them into three immune classes following the manual classification criteria. Since the grids correspond to tumor or tumor-immune coexistence regions, we did not expect grids with a predominance of immune cells. Each grid was categorized into an immune class following the categorical immune cut-offs described above. For WSI representativity, the ratio of grids per immune class was calculated, defining the ratio for the number of selected grids and ROIs per immune class. Then, to optimize the selection of grids, we utilized cosine similarity with a reference set of 72 grids across 12 WSIs. This reference set was built by creating a 3000 x 3000 pixels^2^ grid around each manual ROI. We used PLIP^35^ as the GFM for feature extraction in all reference grids, generating a reference embedding space for each immune class. Then, for grid selection per WSI per immune class, we employed a Hybrid.v2 framework that integrates complementary signals from the GFM (PLIP) and the STM (ASTROS). Specifically, PLIP was applied to extract features from each grid, and a GFM score S_GFM_ was computed as the maximum cosine similarity between the candidate grid embedding and the reference embedding space of the corresponding immune class. Concurrently, ASTROS was applied to every grid, yielding a STM score S_AST_ defined as the maximum detection confidence returned for that grid. The two scores were combined using a data-driven linear weighting scheme in which the relative contribution of each modality was determined by the discriminative spread quantified as the coefficient of variation (CV = σ/μ) of the score’s distribution across the grid population of the current WSI:

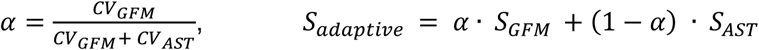

This formulation assigns greater weight to the component that exhibits higher relative variability across grids on a given slide, allowing the pipeline to adapt to slides in which one signal is more informative without requiring manual hyperparameter tuning. Stratified by immune category, the grids selected for each WSI were those that maximized S_adaptive_.

### Automated ROI selection with ASTROS

The selected grids were further analyzed to automate the selection of ROIs with ASTROS, our AI-based Specialist Task-oriented model for ROI selection. ASTROS’s backbone is YOLOv8^52^; we previously compared YOLOv8 with other architectures^53^, showing its superior performance. For model training and cross-validation, a baseline model incorporated the manual ROIs (N = 1,116) from mIF images. Further, to avoid a single-pathologist bias and incorporate pathologists’ heterogeneity, a fine-tuned model included an additional 222 manual ROIs from two independent pathologists following the same selection guidelines. Model training was done with 606 ROIs (58 cases, 34 carcinoma, 17 sarcoma, 6 melanoma, 1 other), using a batch size of 64 and an input resolution of 1280 pixels.

Training was conducted at a learning rate of 0.01, momentum of 0.937, and a weight decay of 0.0005 for up to 1000 epochs, employing an early stopping criterion with patience of 200 epochs. Model performance during training was evaluated using 5-fold cross-validation with a 90/10 train-validation split across 58 slides (35 carcinoma, 6 melanoma, 16 sarcoma, 1 other). ASTROS achieved training losses of 0.523 for bounding box regression, 0.399 for classification, and 1.001 for distribution of focal loss. The Validation test set consisted of 119 slides (21 carcinoma, 31 melanoma, 57 sarcoma, 10 other), and the corresponding losses were 1.686 (box), 0.985 (classification), and 1.510 (distribution focal loss).

At inference, ASTROS was applied to each selected 3,000×3,000-pixel^2^ grid tile, which was internally downsampled to 1,280×1,280 pixels^2^ by the Ultralytics inference engine before prediction. The model localized the highest-confidence immune-infiltrated region within each tile and returned bounding box coordinates, which the engine automatically rescaled back to the native 3,000×3,000-pixel space. The centroid of this predicted bounding box was then used to extract a final ROI of 1,649×1,961 pixels^2^ centered on the detection, matching the dimensions of the manual pathologist ROIs used for training and downstream DSP analysis. This approach allowed ASTROS to be trained efficiently at 1,280 pixels.

For automated mIF-ROI selection, ASTROS analyzed each selected grid, generating ROIs with the highest confidence scores at an IOU threshold of 0.5. For performance evaluation, we calculated the overlap between the automated and manually selected ROIs in two ways: (i) using the estimated grid where the manual ROI was annotated as input, and (ii) using the representativity-optimized grid selected during stage 2 as input. For the evaluation scenario (i), we quantified the mean average precision (mAP) and the intersection over union (IOU) scores between the predictions and the pathologists’ ROI selections.

Two technical caveats. First, to avoid making conclusions from the evaluation scenario (i) that could be reflected by the spatial limitations of the ROI positioning, *i.e.*, a high performance as given due to the limited space within a grid, we simulated a scenario initialized with a target ROI (512 × 512 mm^2^) in a grid of 1.2 × 1.2 mm^2^ over which we randomly placed a second ROI (512 × 512 mm^2^). At each simulated step (N = 1,000), we calculated IOU, Dice score, precision, and recall. The resulting distribution served as a null model reference for randomly placing an ROI, entirely agnostic of tissue properties and limited only by space. Second, to evaluate whether, besides the spatial overlap, the manual and predicted ROIs selected comparable patterns, we quantified and compared the single-cell immune score (scIS) in manual and automated ROIs using a non-parametric Wilcoxon test.

### Comparative performance of generalist and specialist integration

To evaluate the relative contribution of each scoring component of the pipeline, we implemented six experimental arms. The two first arms provide a slide-level population reference against which all AI arms are evaluated. The next four arms, AI arms, differ in their grid scoring function, i.e., the mechanism by which candidate 3,000×3,000 pixel^2^ grids are ranked, and the final six ROIs selected. All AI arms share identical WSI pre-processing, tile extraction, background filtering, and downstream CD45% quantification steps based on our single-cell immune score.

Arm 1: Manual ROIs, pathologist selected manual ROIs per slide based on the immune infiltration patterns serving as the gold standard for ROI level calculations.

<u>Arm 2</u>: WSI reference, systematic calculation of slide-level scIS reference. In this arm, grids ≥80% tissue coverage were retained, scIS calculated at grid level and averaged to produce a WSI-level immune score.

<u>Arm 3</u>: Hybrid.v2, adaptive generalist-specialist integration. Grids were scored by S_adaptive_, integrating both GFM and ASTROS with per-slide data-driven weights. Final ROIs were extracted using ASTROS bounding-box detection on the adaptively selected grids. This arm represented our primary approach by leveraging complementary information across models.

<u>Arm 4</u>: Hybrid.v1, generalist-specialist in tandem. Grids were scored exclusively by their GFM-derived cosine similarity to the reference embedding space (S_GFM_). Final ROIs were extracted using ASTROS bounding-box detection on each selected grid. This arm therefore a sequential formulation more sensitive of the reference grid set, in which semantic GFM guidance governs grid selection while ASTROS performs the final ROI localization.

<u>Arm 5</u>: GFM-only, grids were scored exclusively by their PLIP derived cosine similarity to the reference embedding space (S_GFM_), without any contribution from ASTROS, the specialist model. Final ROIs were obtained by extracting a fixed 1,649×1,961 pixel^2^ crop centered in the selected grid. This arm was designed to assess the sufficiency of GFM-based semantic tissue context guidance to reproduce manual ROI-based and WSI immune scores.

<u>Arm 6</u>: ASTROS-only, systematic extraction of 3,000 × 3,000 pixel^2^ grid from each WSI and directly passed through the ASTROS. Then, each grid received a score equal to the maximum detection confidence returned (S_STM_). Grids were then ranked by S_STM_ in descending order, and the six highest confidence grids were selected. Final ROIs were extracted directly from these selected grids using the bounding box returned by the same ASTROS inference pass. GFM-based feature extraction was not performed in this branch. This arm assessed the contribution of ASTROS in isolation.

To evaluate arm-level differences in CD45% immune infiltration, the distribution of mean CD45% values computed across the six selected ROIs per slide was characterized for all arms and compared using two-sided Wilcoxon rank-sum tests performed at the slide level, with each slide contributing one observation per arm equal to its mean CD45% across the six selected ROIs. Statistical comparisons were conducted at the slide level rather than the tile level to avoid inflation of test statistics driven by the substantially larger number of observations in the WSI Grid and Bootstrap arms relative to the ROI-level arms. All tests were two-sided and a significance threshold of p < 0.05 was applied throughout.

To evaluate agreement with pathologist ROI placement, the Pearson correlation coefficient was computed between each arm’s slide-level mean CD45% estimate and the corresponding manual ROI mean CD45% estimate across all slides. Within-slide consistency was additionally quantified as the inverse of the standard deviation of CD45% estimates across the six selected ROIs per slide, averaged across slides, where higher values indicate more reproducible sampling. Spatial overlap between AI-selected and pathologist-selected ROI positions was quantified using the Hungarian algorithm to find the optimal one-to-one assignment between the six AI and six manual ROIs per slide, with pairwise intersection-over-union (IoU) as the assignment cost; the mean IoU of the optimal assignment was then averaged across slides to yield a per-arm spatial agreement score. The bootstrap percentile of each arm’s mean CD45% estimate was computed by expressing it relative to a null distribution of 1,000 random draws of six tiles per slide from the full WSI-grid tile pool, providing a chance-corrected measure of immune enrichment independent of absolute CD45% levels.

To quantify the computational cost of the GFM component and evaluate whether it is justified by a commensurate gain in accuracy, wall-clock processing time was recorded at component level for each of the slides in both the Hybrid.v2 and ASTROS-only arms, decomposing total per-slide runtime into five mutually exclusive intervals: tile extraction, PLIP embedding (tPLIP), YOLO grid scoring, score combination and selection, and all remaining steps including ROI extraction and re-stitching. The relationship between t_PLIP_ and slide complexity was characterized by Pearson linear regression of t_PLIP_ against the number of candidate grids per slide. The relative PLIP overhead defined as t_PLIP_ as a percentage of total ASTROS-only runtime was summarized by slide complexity group and reported as mean ± standard deviation.

To jointly evaluate accuracy and computational cost, a per-slide efficiency ratio was defined as (MAE_adaptive_ / t_total,adaptive_) / (MAE_ASTROS_ / t_total,ASTROS_), where MAE was computed at n = 6 ROIs per slide against the WSI-grid census reference and values below 1.0 indicate that the Hybrid.v2 arm achieves greater accuracy per unit of compute. Efficiency ratios were computed for all slides with valid MAE and timing data in both arms, summarized by complexity group, and the distribution was tested against the null value of 1.0 using a one-sample Wilcoxon signed-rank test. The per-slide total overhead of the Hybrid.v2 arm relative to ASTROS-only was additionally expressed as a percentage and summarized by complexity group to contextualize the efficiency ratio in terms of absolute time cost.

### Sample size sensitivity

To assess the sensitivity of model performance to sample size (number of ROIs), we conducted a two-level sampling analysis consistent with classical sampling theory. For within-slide level analysis, we quantified how the precision of the slide-level immune score estimation depends on the number of ROIs sampled. Using the Hybrid.v2 arm ROIs (n=6 per slide), we computed the within-slide standard deviation of the mean immune score (mIF-scIS, CD45%) across all subsets of size n = {1, …, 5}, averaging over all slides. A power-law model (σ∝n^a^) was fitted to the mean standard deviation as a function of sample size using nonlinear least squares. The theoretical 1/√n decay (α = −0.50) was overlaid as a reference. Mean Absolute Error (MAE) against the pathologist-annotated manual mean was similarly computed at each n, and the proportion of total achievable MAE reduction captured at each n was derived relative to the asymptotic value at n=6 using point percentage as a metric.

For a cohort-level bootstrap, we performed a bootstrap subsampling analysis over the full pool of 1,036 Hybrid.v2 ROIs. For each target cohort size N (ranging from N=5 to N=1,036 in logarithmically spaced steps), we drew 5,000 independent random samples of N ROIs without replacement and computed six performance metrics against both the manual pathologist reference and the WSI grid reference: Pearson’s r, Lin’s concordance correlation coefficient (CCC), MAE, and systematic bias (mean signed error). For each N, the mean point estimate and 95% confidence interval (2.5^th^–97.5^th^ percentile of the bootstrap distribution) were recorded. The 95%CI width as a function of N was fit with a power law model (width ∝ N^α^) on a log-log scale using ordinary least squares regression, and the empirical scaling exponent was compared to the theoretical −0.50 expected under independent sampling.

### Virtual support with virtual mIF automated ROI selection

To further enhance the versatility of our approach, we created a virtual support framework that enabled preliminary visualization of ROI selection without requiring mIF as a step in the workflow. For this purpose, the mIF sections used in the DSP assay were retrieved, washed, and re-stained with H&E, followed by digitization, same-tissue section H&E-to-mIF image alignment, and quantitative assessment of the registration. The virtual support stage is triggered by an input H&E that, during the processing, was virtually stained for morphological markers, followed by our automated ROI selection process, resulting in a virtual mIF (vmIF) with selected ROIs.

### Restaining mIF with H&E for image registration

The restained H&E images were digitized at 40× magnification, resolution = 0.251 μm/pixel (Aperio AT2 scanner platform, Leica Biosystems, Wetzlar, Germany) and registered to their paired mIF (40× magnification, resolution = 0.4002). For image registration, we applied sequentially rigid and affine registration using wsireg (https://github.com/NHPatterson/wsireg), registering the H&E image to the mIF. We assessed the registration performance using the Dice and multi-scale structural similarity (MS-SSIM) scores^54^ (Python, scikit-image) applied to binary nuclear masks obtained through cell identification in QuPath^55^. We used similar hyperparameters for nuclear detection in both modalities to minimize the confounding effect of nuclear detection models (**Supplementary Table 5**). Since the cell nucleus is a common feature between mIF and H&E assays, we used nuclear segmentation as a more robust approach than cell segmentation to the variability between both modalities. Although nuclear segmentation may include false positives or hypersegmentation, we only utilized this detection to assess the registration performance.

### Virtual mIF from H&E

With 168 cases registered, we trained Pix2Pix^41^, a conditional generative adversarial network (GAN) that allowed the virtual mIF staining of H&E images. Pix2Pix utilized a paired set of images and consisted of two main architectural components: a generator and a discriminator. The generator used H&E and mIF images to learn plausible pairs for the modality translation. The discriminator learned to differentiate the most plausible translation by comparing it with the target image (mIF).

For our dataset, Pix2Pix was trained on 147,492 mIF/H&E paired tiles stratified across 28 WSI (14 Carcinoma, 7 Melanoma, 7 Sarcoma). For validation, 52,040 paired tiles were included, stratified across 10 samples (6 Carcinoma, 2 Melanoma, 2 Sarcoma). A standard Pix2Pix architecture was implemented with a generator (netG) as a U-Net (256×256 receptive fields) and the discriminator utilizing a PatchGan. Both networks employed batch normalization throughout. Input images were preprocessed using a resize and crop strategy, both set to 256 pixels. The model was then optimized using a Vanilla GAN objective combined with a pixel-wise L1 reconstruction loss weight by λ = 100 to encourage structural fidelity between input and output. Standard optimization was applied using the Adam optimizer, with a learning rate of 0.0002, β₁ = 0.5, and linear learning rate decay post 100 epochs. Training was performed for 1,000 epochs, and at the final training iteration, the model exhibited loss values of 6.36 for generator L1 loss, 2.01 for generator adversarial loss, 0.67 for discriminator loss (real mIF images), and 0.47 for discriminator loss (virtual mIF images), indicating stable convergence and effective adversarial learning. To quantify the virtual staining performance, we followed a similar approach to the registration evaluation, using quantitative image metrics for image quality assessment (MS-SSIM) and nuclear mask overlap (Dice score).

### ASTROS ROIs selection on synthetic mIF and comparison with mIF

For automated ROI selection in the generated vmIF, we followed the same protocol used for mIF with our hybrid generalist-specialist workflow using Hybrid.v2. We quantified the performance by calculating pairwise IOU scores between vmIF and mIF ROIs. Furthermore, for each WSI, we average the single-cell immune scores (scIS, %CD45^+^ cells) and compare them between the mIF-ROIs and vmIF ROIs suing a Wilcoxon test and rank-biserial correlation r for power calculation. Finally, as the correlation between immune scores in the automated ROIs selected in mIFs and vmIFs can be influenced by registration and virtual staining performance, we included the corresponding Dice scores as covariates for the partial correlation between mIF-scIS and vmIF-scIS, comparing them with the correlation without accounting for covariates (R package: ppcor).

### ROI selection for Visium platforms Data description

An in-house dataset, Visium cohort, was utilized to demonstrate the framework’s flexibility in integrating with spatial transcriptomic platforms. We included 66 candidate samples for ST analysis with 10x’s Visium retrospectively annotated (cholangiocarcinoma-CCA, lung adenocarcinoma-LUNG) and Visium HD (glioblastoma-GBM, upper tract urothelial carcinoma-UTUC, cholangiocarcinoma-CCA). We included four CCA cases as a proof of concept for the prospective application of AI-assisted ROI selection. As specified by 10x Genomics, for each Visium assay, an H&E tissue section was prepared and digitized (resolution = 0.2513 mpp). Additionally, an ROI was manually selected in each tissue section (Visium: 11×11 mm^2^ and Visium HD: 6.5×6.5 mm^2^) to perform image-based transcript capture. The criteria for pathologists’ manual ROI selection varied across tissues, including representativity, intratumor heterogeneity, or the detection of specific morphological patterns.

### Scenarios for ROI selection in the generalist-specialist framework

To illustrate some of the potential use cases, we develop three scenarios that demonstrate how GFM and STM can be used standalone or both integrated with a pathologist’s input to give a streamlined solution to common needs. Although these represent some possible uses, the expanding modularity can accommodate a different number of models tailored to other applications and the specific goals of a study.

Unless otherwise stated, we used UNI^29^ as the GFM to extract embeddings at the tile level. For the STM model, we used a pretrained model for tissue segmentation (TMESegformer^42^). Due to the pathologist’s specific requirements, scenario three was only used for CCA, including coarse manual annotations. A sliding window approach was applied in each case, with a size equal to the manual ROI and a stride of 12 pixels.

<u>Scenario 1: capture representative tumor areas:</u> The goal of this scenario was to showcase GFM usage for selecting ROIs based on a set of manually chosen ROIs by training a minimalist morphology-based ROI selection model that learns the embedding-space signature of manually selected ROIs. This was to fulfill a common need to select ROIs capturing representative tumor areas while maximizing viable tissue coverage within the ROI.

To develop this scenario, the model was trained with ROIs manually selected for LUNG, CCA, and UTUC. Although diverse in morphology, this set shared a minimal set of features, such as viable tumor tissue. The tissue was segmented and tiled (256 × 256 pixels^2^) to extract tile-level embeddings. For each tile, a binary label was assigned based on whether its center was within the manual ROI (positive) or fell outside (negative). To compensate for imbalanced datasets, negative non-ROI tiles are randomly downsampled to match the number of positive ROI tiles. To train the ROI-tile model, embeddings were normalized using StandardScaler (Python: scikit-learn), and a Logistic Regression model was trained to classify ROI vs non-ROI tiles. An 80/20 train-validation split is applied for evaluation. During training, the primary evaluation metric was the area under the receiver operator curve (AUC). When evaluated on the independent validation set, the model achieved an area under the ROC curve (AUC) of 0.963, reflecting excellent discriminative ability. The ROC profile indicated that the classifier maintained high sensitivity across a broad range of specificity levels, suggesting that the combination of balanced sampling, standardized features, and a regularized logistic regression model produced a robust and reliable predictor.

To assess the scenario, we employed a diverse set of metrics to both visually and quantitatively evaluate the prediction. Each tile received a positive-classification probability, resulting in a probability map over the entire slide, capturing the spatial distribution of “ROI-likeness”. To reach an optimized ROI selection, a rectangular window matching the size of the training ROIs was slid over the WSI. At each iteration, the average ROI probability of the tiles inside the moving window was calculated. The window with the highest average ROI probability was selected as the predicted ROI, identifying the most viable region in terms of morphology. The tile probabilities were visualized as a heatmap to provide an intuitive, spatial overview of where ROI-like morphology is concentrated. Finally, we calculated the intersection over union score (IOU) to measure the overlap between the predicted ROI and the manual annotation.

<u>Scenario 2: Targeted inclusion of certain tissue compartments:</u> In this scenario, we addressed specific requirements for including or excluding certain tissue components, such as necrosis. For this purpose, a specialist TMESegformer model was used for TME segmentation and optimization of the tumor-to-necrosis ratio in ROIs, identifying viable regions suitable for spatial transcriptomics while harnessing the performance and explainability of STMs.

In this case, each WSI was processed using the TMESegformer^42^ model pre-trained on lung adenocarcinoma. Hence, we applied it to the lung cases in our cohort. In short, the model segmented each WSI into 10 tissue classes: tumor, stroma, immune (macrophage/inflammatory), necrosis, alveoli, bronchi epithelium, vessels, adipose tissue, and muscle. The predicted segmentation was saved as a dense mask. To systematically sample the tissue, a rectangular ROI window, with a size equal to the manual ROI, was slid across the TMESegformer prediction mask. For each window, the proportion of pixels belonging to each TME class was computed. A composite score was calculated as the ratio between tumor and necrosis. The window with the highest composite TME score was selected as the ROI. For evaluation, the IOU score was calculated between the manual and predicted ROIs. Pearson’s correlation was used to assess the association between tumor and necrosis percentages of manual and predicted ROIs.

<u>Scenario 3: Human-AI collaboration:</u> A final scenario entailed the interaction of GFM, STM, and the pathologist’s input in the form of a manually segmented mask. This scenario can be useful in cases when ROI needs to be directed to a specific region of the tissue, coarsely defined by manual input (*e.g.,* tumor bed). We aimed to design a cascaded pipeline that used pathologist-provided tumor masks to guide ROI selection, enabling fine-grained detection of tissue viability. The method integrated expert annotations to anchor the model to the tumor region, an STM (TMESegformer) for TME segmentation within the pathologist-defined tumor mask, and a GFM capturing subtle and representative morphological patterns for final ROI selection.

Two pathologists (KP, CE) annotated a consensus tumor region in each WSI using QuPath^55^. Then, this region was intersected with the TMESegformer prediction mask. Tissue components, including necrosis, tumor, immune, and stromal components, were identified at the pixel level. A filter with attention to the tumor and stromal compartments was applied before deploying the GFM on this tissue compartment. The embeddings captured subtle morphological differences in the tumor compartment, enabling fine clustering or classification. For ROI selection optimization from a moving window, the region with the highest embedding cosine similarity with the pathologist’s annotated tumor bed was selected, ensuring morphological representativity of the WSI with attention to the coarse pathologist’s annotation. The performance of ROI prediction was evaluated by calculating the IOU score between the manually annotated and predicted ROIs.

### Prospective ROI selection

As proof of concept of the implementation, we processed four cases of cholangiocarcinoma for Visium HD (6.5×6.5 mm^2^ ROI). We applied scenarios I (only GFM) and III with minimal manual input in the form of coarse tumor bed annotation. Each WSI with the proposed ROIs, from scenarios I and III, was presented to a pathologist (CE). The pathologist decided if one of the ROIs selected an appropriate region, avoiding artifacts, necrosis, or hemorrhages as much as possible. An option of rejecting the proposed ROIs was also presented. After the assessment and decision, the ROI was transferred to the physical slide, completing the ROI selection for Visium HD.

## DATA AVAILABILITY

Data has been deposited in Zenodo (https://zenodo.org/records/19389837) and full access can be requested from the corresponding author (Y.Y.) via Zenodo. De-identified clinical data is also available under restricted access due to patient privacy policies.

## CODE AVAILABILITY

This work uses publicly available models referenced throughout the manuscript. The algorithm is described at a functional level in the Methods section. Full implementation details will be made available upon a reasonable request.

## ACKNOWLEDGMENTS

This work is supported by the Spatial Ecology & QUantitative pathOlogy Image Analytical platform (SEQUOIA) through the MD Anderson STrategic Research Initiative Development Program (STRIDE) and Patient Mosaic^TM^ project at The University of Texas MD Anderson Cancer Center.

## AUTHOR CONTRIBUTIONS

Y.Y. and S.P.C. had the original idea. S.P.C., T.G., Y.Y. made work conceptualization. Patient Mosaic, L.S-S., A.S., B.L.R., J.T.H., C.E., X.P., L.K., A.L., S.P.C. participated in data acquisition and curation. T.G., S.P.C., P.C., P.A., and Y.S. performed data processing and coding. S.P.C. and T.G. did formal analysis. L.S-S., A.S., K.P., M.E.S., J.T.H., and C.E. did pathology assessment. Project administration was done by Y.Y., B.L.R., L.S-S., Patient Mosaic. S.P.C. and Y.Y. jointly supervised this study. S.P.C. and T.G. drafted the initial manuscript draft. All coauthors participated in reviewing the final manuscript.

