## Supplementary for "A modular generalist-specialist AI framework for ROI selection across spatial profiling workflow"

### TITLE

### Supplementary Figures

Supplementary Fig. 1 Agreement between manual ROI and whole-slide image (WSI) immune scores.

Supplementary Fig. 2 Sample size sensitivity analysis of Hybrid.v2 ROI-based CD45% estimation across within-slide and cohort-level scales.

Supplementary Fig. 3 Comparison of biological performance between four methods for ROI selection.

Supplementary Fig. 4 Comparison of computing performance between four methods for ROI selection.

Supplementary Fig. 5 Specialist Task-oriented TMESegformer enables ROI selection for Visium, minimizing non-viable areas.

### Supplementary Tables

Supplementary Table 1 Patient Mosaic cohort composition.

Supplementary Table 2 Morphological panel for immunofluorescence.

Supplementary Table 3 Target proteins for the DSP spatial proteomic assay.

Supplementary Table 4 Overall variability metrics for marker expression across tissue compartments.

Supplementary Table 5 Category-stratified variability metrics for marker expression across tissue compartments.

Supplementary Table 6 Performance of ROI selection at manually predefined input grids.

Supplementary Table 7 Nuclear segmentation parameters used in QuPath to evaluate registration.

**Annex I**: Patient Mosaic Team

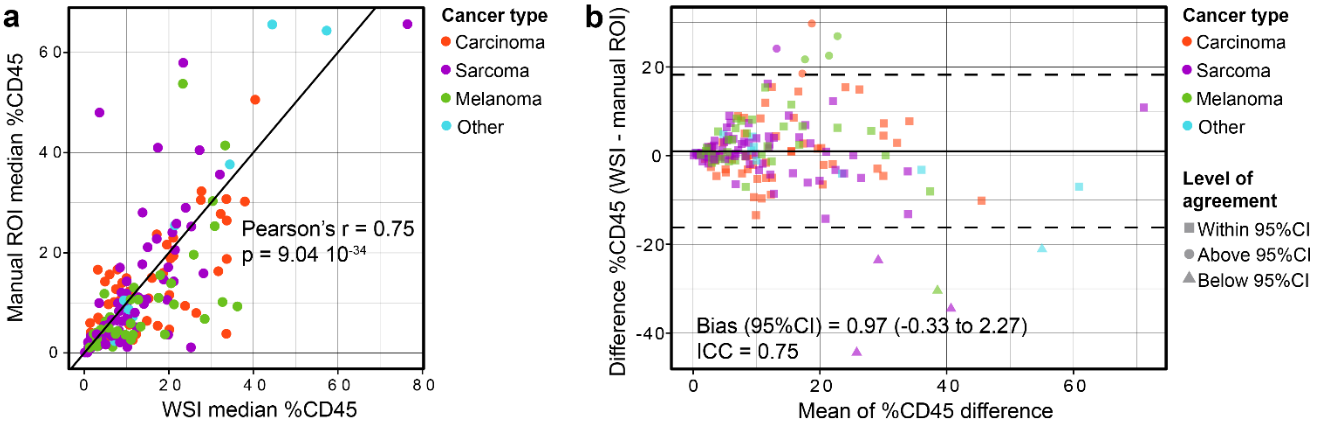


**Supplementary Fig. 1** Agreement between manual ROI and whole-slide image (WSI) immune scores. **a** Correlation between automated immune scores from manual ROIs and WSIs. Each dot represents the median value per sample. **b** Bland-Altman analysis of the agreement between ROI and WSI immune scores. The plot shows the difference between paired median CD45% measurements versus their mean (n = 178). The central solid line denotes the mean bias, and dashed lines denote the 95% limits of agreement. The estimated bias was 0.97, with limits of agreement from -16.25 to 18.19. Point color corresponds to cancer type, and in (**b**), shape corresponds to the position relative to the limits of agreement.

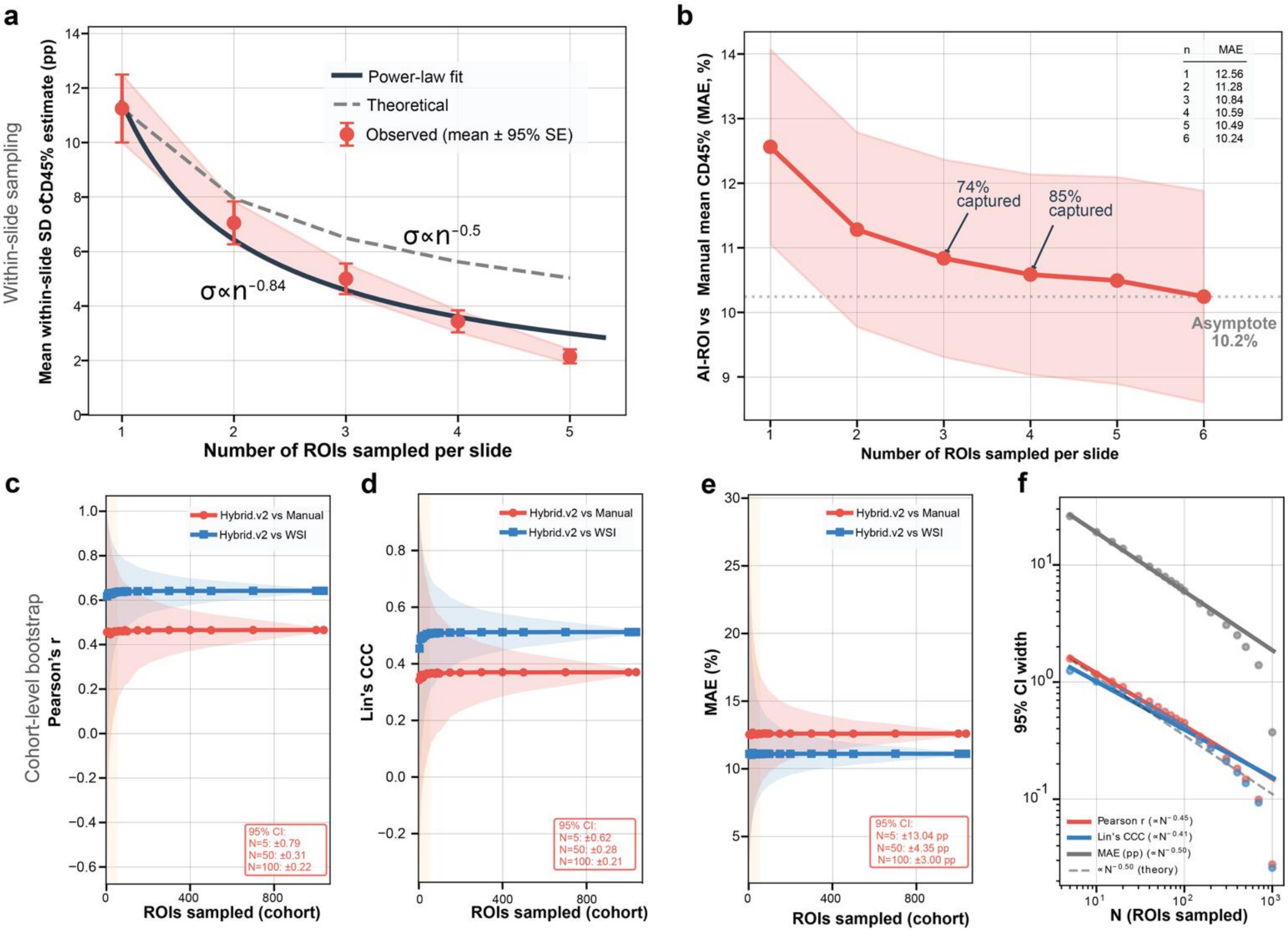


**Supplementary Fig. 2** Sample size sensitivity analysis of Hybrid.v2 ROI-based CD45% estimation across within-slide and cohort-level scales. **a** Within-slide estimation precision as a function of the number of ROIs sampled per slide. The mean within-slide standard deviation of the CD45% estimate (±95% SE) decays as $\sigma \propto n^{-0.84}$ (dark line), converging faster than the theoretical $1/\sqrt{n}$ relationship (dashed grey). **b** Mean Absolute Error (MAE) vs manual pathologist CD45% mean as a function of n ROIs per slide (±95% SE). **c-e** Cohort-level bootstrap subsampling analysis (5,000 iterations per sample size) over the full pool of 1,036 Hyrbid.v2 ROIs. Performance point estimates for **(c)** Pearson's r, **(d)** Lin's concordance correlation coefficient (CCC), and **(e)** mean absolute error (MAE) are shown against both manual pathologist (red) and WSI grid references (blue), with 95% bootstrap CI ribbons. Orange shading denotes the high-uncertainty zone (N < 50). Dashed vertical lines mark N=50 (orange) and N=100 (grey). **f** Log-log plot of 95% CI width as a function of N for Pearson's r, Lin's CCC, and MAE.

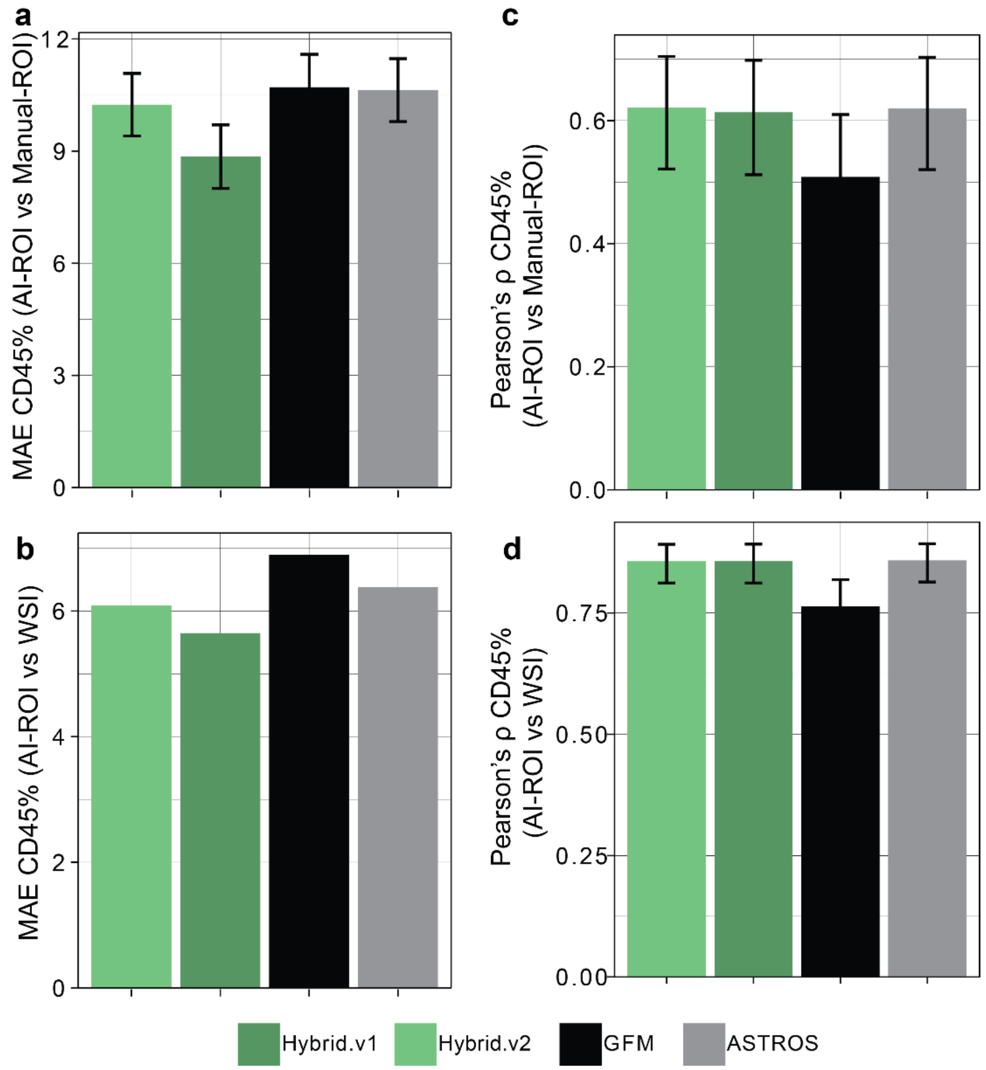


**Supplementary Fig. 3** Comparison of biological performance between four methods for ROI selection. **a-b** Bar plot of mean absolute error (MAE) between AI-ROIs CD45% scIS and (**a**) manual ROIs (± 95%CI) and (**b**) WSI-level estimates. **c-d** Bars showing the Pearson's coefficient (± 95%CI) for the correlation between AI-based ROIs scIS and (**c**) manual ROIs scIS and (**d**) WSI-level scIS.

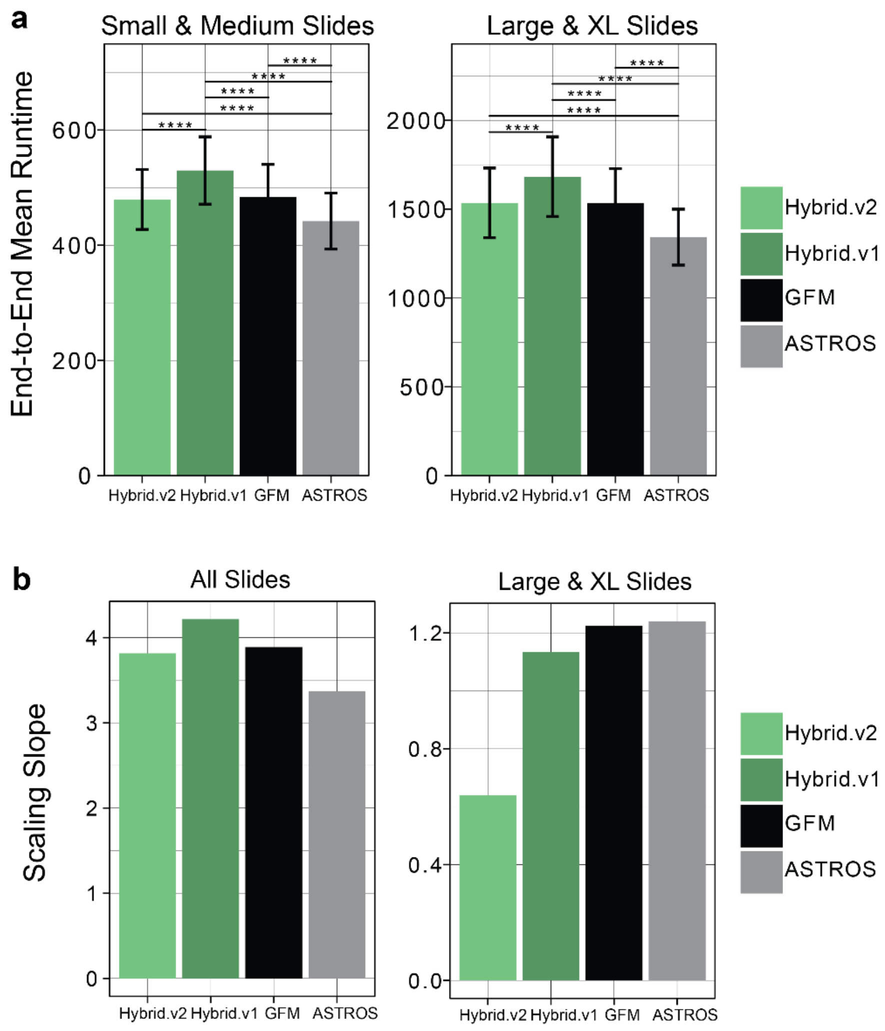


**Supplementary Fig. 4** Comparison of computing performance between four methods for ROI selection. **a** End-to-end runtime calculated as the wall-clock time from pipeline entry to final ROI selection. Bars show slide-count–weighted mean runtime for each method within the pooled Small/Medium and Large/XL groups; error bars indicate 95% confidence intervals. Statistical comparisons were performed only within each pooled size stratum, with Bonferroni-Holm adjusted p values shown ****p<0.0001. **b** Bars showing the value for the scaling slope (seconds/tile) quantified as the change in runtime per additional tile in one slide.

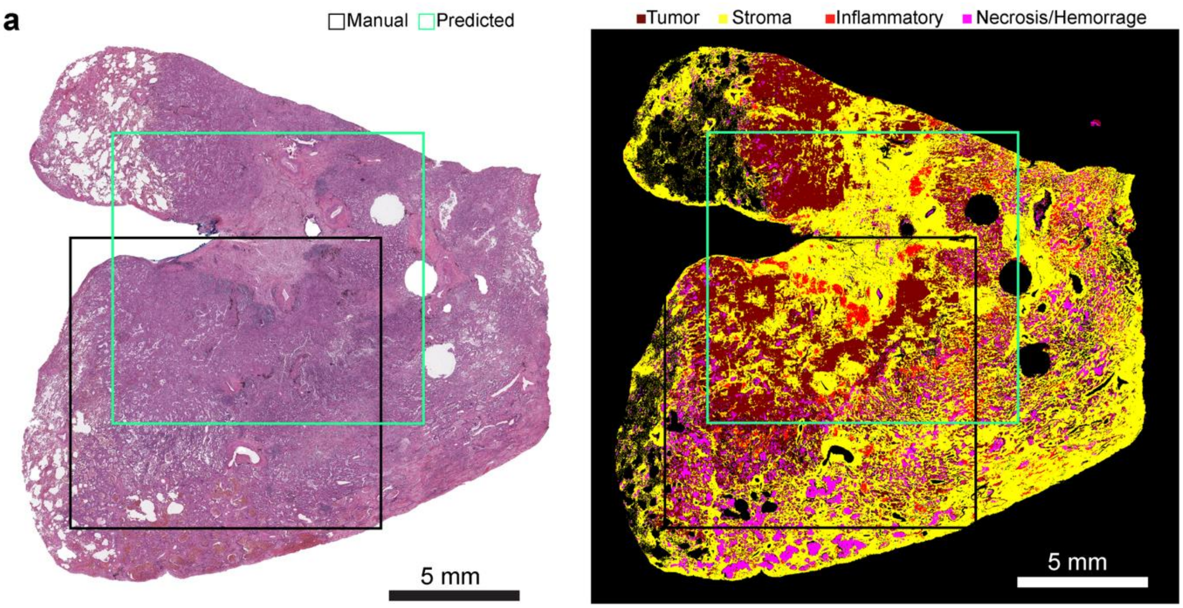


**Supplementary Fig. 5** Specialist Task-oriented TMESegformer enables ROI selection for Visium minimizing non-viable areas. **a** A case for ROI prediction through the application of an STM to minimize necrosis, hemorrhages, and maximize tumor content. A lower intersection between the manual and predicted ROIs (IOU = 0.34) can be attributed to the strict optimization algorithm minimizing non-viable tissue, such as hemorrhage or necrosis.

**Supplementary Table 1.** Patient Mosaic DSP cohort composition. Regions of interest (ROI). The average number of ROIs per sample is rounded to the nearest integer.

| Major tumor type | Diagnosis | Number of cases | Number of ROIs Total (average per sample) | Automated segmentation strategy |
|---|---|---|---|---|
| Carcinoma | Papillary urothelial carcinoma | 22 | 134 (6) | Tumor/TME |
| | Cholangiocarcinoma | 6 | 43 (7) | Tumor/TME |
| | Colorectal adenocarcinoma | 5 | 36 (7) | Tumor/TME |
| | Mucoepidermoid carcinoma | 2 | 15 (8) | Tumor/TME |
| | Esophageal adenocarcinoma | 2 | 11 (6) | Tumor/TME |
| | Urothelial carcinoma | 2 | 11 (6) | Tumor/TME |
| | Penile squamous cell carcinoma | 2 | 10 (5) | Tumor/TME |
| | Rectal squamous cell carcinoma | 2 | 10 (5) | Tumor/TME |
| | Neuroendocrine carcinoma | 1 | 9 (9) | Tumor/TME |
| | Papillary thyroid carcinoma | 1 | 9 (9) | Tumor/TME |
| | Hepatocellular carcinoma | 1 | 6 (6) | Tumor/TME |
| | Hurtle cell carcinoma | 1 | 6 (6) | Tumor/TME |
| | Lung adenocarcinoma | 1 | 6 (6) | Tumor/TME |
| | Ovarian clear cell carcinoma | 2 | 12 (6) | Tumor/TME |
| | Ovarian serous carcinoma | 1 | 6 (6) | Tumor/TME |
| | Renal cell carcinoma | 1 | 6 (6) | Tumor/TME |
| | Appendiceal adenocarcinoma | 1 | 4 (4) | Tumor/TME |
| | Undifferentiated thyroid carcinoma | 1 | 6 (6) | Tumor/TME |
| | Medullary thyroid carcinoma | 1 | 6 (6) | Tumor |
| Melanoma | Anorectal melanoma | 11 | 85 (8) | Tumor/TME |
| | Vulvovaginal melanoma | 6 | 43 (7) | Tumor/TME |
| | Uveal melanoma | 2 | 17 (8) | Tumor/TME |
| | Sinonasal melanoma | 7 | 44 (6) | Tumor/TME |
| | Conjunctivital melanoma | 1 | 6 (6) | Tumor/TME |
| | Cutaneous melanoma | 1 | 6 (6) | Tumor/TME |
| | Nasolacrimal melanoma | 1 | 6 (6) | Tumor/TME |
| Sarcoma | Alveolar soft part sarcoma | 1 | 3 (3) | Tumor |
| | Chondrosarcoma | 7 | 62 (9) | Tumor/TME |
| | Dedifferentiated liposarcoma | 7 | 60 (9) | Tumor |
| | Dermatofibrosarcoma | 1 | 6 (6) | Tumor |
| | Desmoplastic small round cell tumor | 1 | 17 (17) | Tumor/TME |
| | Extra skeletal chondrosarcoma | 1 | 12 (12) | Tumor |
| | Low-grade fibromyxoid sarcoma | 1 | 6 (6) | Tumor |
| | Leiomyosarcoma | 11 | 81 (7) | Tumor |
| | Liposarcoma | 4 | 40 (10) | Tumor |
| | Malignant sarcomatoid tumor | 1 | 6 (6) | Tumor |
| | Osteosarcoma | 1 | 6 (6) | Tumor |
| | Pleomorphic sarcoma | 2 | 12 (6) | Tumor |
| | Rhabdomyosarcoma | 1 | 6 (6) | Tumor |
| | Spindle cell sarcoma | 5 | 31 (6) | Tumor |

| | | | | |
|---|---|---|---|---|
| | Synovial sarcoma | 7 | 47 (7) | Tumor |
| | Unclassified small round cell sarcoma | 1 | 6 (6) | Tumor |
| | Undifferentiated pleomorphic sarcoma | 6 | 30 (5) | Tumor |
| | Uterine adenosarcoma | 2 | 18 (9) | Tumor |
| | Well-differentiated liposarcoma | 7 | 48 (7) | Tumor |
| Other | Chordoma | 1 | 4 (4) | Tumor/TME |
| | Invasive thymoma | 1 | 6 (6) | Tumor/TME |
| | Low-grade sarcomatoid process | 1 | 6 (6) | Tumor |
| | Mesothelioma | 2 | 17 (8) | Tumor/TME |
| | Ossifying fibromyxoid tumor | 1 | 6 (6) | Tumor |
| | Spindle cell neoplasm | 1 | 6 (6) | Tumor |
| | Uterine carcinosarcoma | 1 | 9 (9) | Tumor/TME |
| | Well-differentiated papillary mesothelial tumor | 1 | 6 (6) | Tumor/TME |
| | Pancreas solid pseudopapillary neoplasm | 1 | 6 (6) | Tumor/TME |

**Supplementary Table 2.** Morphological panel for visualization of immunofluorescence during the GeoMx DSP workflow.

| Markers | Target | Clone | Vendor | Catalog # | Labeling Fluorophore | RRID |
|---|---|---|---|---|---|---|
| Syto 13 | Nuclei | N/A | Nanostring | 121300319 | AF488 | |
| PanCK | Tumor cells epithelial | AE1+AE3 | Nanostring/Novus | NBP2-33200 | AF532 | AB_3284598 |
| CD45 | Immune cells | 2B11+PD7/26 | Nanostring/CST | 13917 | AF594 | AB_2797987 |
| S100B | Tumor cells melanoma | S100B/1706R | Nanostring / Novus | NBP2-54426 | AF532 | AB_3335461 |
| Synaptophysin | Tumor cells neuroendocrine | YE269 | abcam | ab 196166 | AF647 | AB_3097754 |

**Supplementary Table 3.** Target proteins for the spatial proteomic assay (NanoString's GeoMx DSP probes).

| Target protein | Gene | Accession | Protein ID | Annotations |
|---|---|---|---|---|
| Beta-2-microglobulin | B2M | P61769 | DPROT_00010.1 | Antigen Presentation, Tumor |
| CD11c | ITGAX | H3BN02, P20702 | DPROT_00011.1 | Dendritic cell, Myeloid cells |
| CD14 | CD14 | P08571 | DPROT_00051.1 | Inflammation, Monocyte, Myeloid |
| CD163 | CD163 | Q86VB7 | DPROT_00052.1 | Macrophage, Microglia, Myeloid |
| CD20 | MS4A1 | A0A024R507, P11836 | DPROT_00012.1 | B cells |
| CD3 | CD3D, CD3G, CD3E | P09693, B0YIY4, B0YIY5, P04234, P07766 | DPROT_00013.1 | T cells |
| CD34 | CD34 | P28906 | DPROT_00048.1 | Hematopoietic |
| CD4 | CD4 | B4DT49, B0AZV7, P01730 | DPROT_00014.1 | Myeloid, T cells, Th cells |
| CD45 | PTPRC | P08575, X6R433, M9MML4, A0A0A0MT22 | DPROT_00015.1 | Inflammation, Microglia, Total Immune |
| CD45RO | PTPRC | P08575, X6R433, M9MML4, A0A0A0MT22 | DPROT_00046.1 | Memory, T cells |
| CD56 | NCAM1 | P13591, A0A087WWD4 | DPROT_00016.1 | NK cells |
| CD66b | CEACAM8 | B4DLI3, P31997, Q0Z7S6 | DPROT_00049.1 | Myeloid, Neutrophil |
| CD68 | CD68 | P34810 | DPROT_00006.1 | M2 Macrophage, Macrophage, Microglia, Myeloid |
| CD8 | CD8A | P01732, Q6ZVS2, Q8TAW8 | DPROT_00017.1 | CD8 T cells, T cells |
| CTLA-4 | CTLA4 | P16410 | DPROT_00395.1 | Checkpoint, T cell Activation, T cells, Th cells |
| FAP-alpha | FAP | Q12884, B4DLR2 | DPROT_00050.1 | Fibroblasts, Stroma |
| Fibronectin | FN1 | Q6MZM7, Q6MZF4, P02751, B7ZLE5, Q9UQS6, Q6N084 | DPROT_00024.1 | Fibroblasts, Stroma |

| | | | | |
|---|---|---|---|---|
| FOXP3 | FOXP3 | Q9BZS1, B7ZLG1 | DPROT_00047.1 | T cells, Th cells, Tregs |
| GZMB | GZMB | P10144, Q67BC3, Q6XGZ4, J3KQ52 | DPROT_00019.1 | Cytotoxicity, T cell Activation |
| HLA-DR | HLA-DRA | A0A0G2JMH6, P01903 | DPROT_00007.1 | Antigen Presentation, MHC2, Microglia |
| Ki-67 | MKI67 | P46013 | DPROT_00009.1 | Proliferation |
| PanCk | KRT6B, KRT16, KRT2, KRT3, KRT19, KRT10, KRT8, KRT6A, KRT1, KRT14, KRT5 | P12035, P13645, P13647, A0A0S2Z428, P35908, P02533, P05787, P04259, P08727, Q7L4M3, P02538, P04264, P08779 | DPROT_00022.1 | Epithelial, Tumor |
| PD-1 | PDCD1 | A0A0M3M0G7, Q15116 | DPROT_00004.1 | Checkpoint, T cell Activation, T cells |
| PD-L1 | CD274 | Q0GN75, Q9NZQ7 | DPROT_00021.1 | Checkpoint, Myeloid Activation |
| SMA | ACTA2 | P62736, D2JYH4 | DPROT_00023.1 | Stroma |
| GAPDH | GAPDH | P04406, V9HVZ4 | DPROT_00020.1 | Housekeepers, Housekeeping |
| Histone H3 | H3C1 | P68431 | DPROT_00005.1 | Housekeepers, Housekeeping |
| S6 | RPS6 | P62753, A2A3R6 | DPROT_00008.1 | Housekeepers, Housekeeping |
| Ms IgG1 | | | DPROT_00002.1 | Background |
| Ms IgG2a | | | DPROT_00003.1 | Background |
| Rb IgG | | | DPROT_00001.1 | Background |

**Supplementary Table 4 Overall variability metrics for marker expression across tissue compartments.** Linear mixed-effects models were fitted for each marker and tissue compartment with tumor type included as a fixed effect and sample as a random intercept. The table reports the between-sample variance, residual variance, and intraclass correlation coefficient (ICC), together with the overall mean, standard deviation (SD), and coefficient of variation (CV) of the measured values. Higher ICC values indicate a greater proportion of total variance attributable to differences between samples. Shading intensity increases with value, with darker cells indicating higher between-sample variance or ICC.

| | | Variance | | | Overall | | |
|---|---|---|---|---|---|---|---|
| Segment | Protein | Sample | Residual | ICC | Mean | SD | CV |
| Tumor | PD-1 | 0.537 | 0.406 | 0.570 | 0.142 | 0.966 | 6.786 |
| Tumor | PD-L1 | 0.596 | 0.289 | 0.673 | -0.055 | 0.967 | -17.549 |
| Tumor | CD45 | 0.558 | 0.283 | 0.663 | -0.216 | 0.926 | -4.284 |
| Tumor | Fibronectin | 0.405 | 0.260 | 0.609 | -0.116 | 0.848 | -7.288 |
| Tumor | Ki-67 | 0.673 | 0.239 | 0.738 | 0.114 | 1.046 | 9.143 |
| Tumor | HLA-DR | 0.600 | 0.319 | 0.653 | -0.115 | 0.987 | -8.562 |
| TME | PD-1 | 0.306 | 0.516 | 0.372 | -0.306 | 1.004 | -3.288 |
| TME | PD-L1 | 0.643 | 0.433 | 0.597 | 0.118 | 1.058 | 8.947 |
| TME | CD45 | 0.604 | 0.368 | 0.622 | 0.464 | 0.997 | 2.149 |
| TME | Fibronectin | 0.455 | 0.239 | 0.655 | 0.250 | 1.231 | 4.928 |
| TME | Ki-67 | 0.458 | 0.236 | 0.660 | -0.246 | 0.843 | -3.435 |
| TME | HLA-DR | 0.582 | 0.390 | 0.599 | 0.248 | 0.983 | 3.971 |

**Supplementary Table 5.** Category-stratified variability metrics for marker expression across tissue compartments. For each marker and tissue compartment, variability was summarized across samples within each category. The table reports the number of samples, mean, standard deviation (SD), interquartile range (IQR), and coefficient of variation (CV) of the sample-level mean values. Cell shading is scaled by magnitude, such that darker shading denotes higher IQR or SD values.

| Segment | Protein | Category | Mean | SD | IQR | CV |
|---|---|---|---:|---:|---:|---:|
| Tumor | PD-1 | Carcinoma | 0.143 | 0.657 | 0.766 | 4.587 |
| Tumor | PD-1 | Melanoma | 0.338 | 0.710 | 0.995 | 2.103 |
| Tumor | PD-1 | Other | 0.336 | 0.730 | 0.977 | 2.172 |
| Tumor | PD-1 | Sarcoma | 0.078 | 0.898 | 1.071 | 11.492 |
| Tumor | PD-L1 | Carcinoma | 0.041 | 0.871 | 0.667 | 21.396 |
| Tumor | PD-L1 | Melanoma | 0.429 | 0.627 | 0.609 | 1.462 |
| Tumor | PD-L1 | Other | 0.212 | 0.702 | 0.661 | 3.312 |
| Tumor | PD-L1 | Sarcoma | -0.263 | 0.851 | 0.964 | -3.239 |
| Tumor | CD45 | Carcinoma | -0.305 | 0.701 | 0.814 | -2.299 |
| Tumor | CD45 | Melanoma | 0.158 | 0.759 | 0.802 | 4.799 |
| Tumor | CD45 | Other | 0.236 | 0.941 | 1.190 | 3.981 |
| Tumor | CD45 | Sarcoma | -0.320 | 0.834 | 0.894 | -2.610 |
| Tumor | Fibronectin | Carcinoma | -0.401 | 0.615 | 0.995 | -1.533 |
| Tumor | Fibronectin | Melanoma | -0.370 | 0.673 | 1.219 | -1.821 |
| Tumor | Fibronectin | Other | 0.328 | 0.963 | 1.470 | 2.931 |
| Tumor | Fibronectin | Sarcoma | 0.173 | 0.681 | 0.793 | 3.936 |
| Tumor | Ki-67 | Carcinoma | 0.433 | 0.838 | 1.145 | 1.933 |
| Tumor | Ki-67 | Melanoma | 0.834 | 0.867 | 0.867 | 1.039 |
| Tumor | Ki-67 | Other | 0.126 | 0.828 | 0.812 | 6.579 |
| Tumor | Ki-67 | Sarcoma | -0.290 | 0.854 | 1.030 | -2.942 |
| Tumor | HLA-DR | Carcinoma | -0.321 | 0.643 | 0.739 | -2.002 |
| Tumor | HLA-DR | Melanoma | 0.467 | 0.906 | 1.117 | 1.939 |
| Tumor | HLA-DR | Other | 0.321 | 0.965 | 1.399 | 3.003 |
| Tumor | HLA-DR | Sarcoma | -0.196 | 0.868 | 1.031 | -4.436 |

| Segment | Protein | Category | Mean | SD | IQR | CV |
|---|---|---|---:|---:|---:|---:|
| TME | PD-1 | Carcinoma | -0.090 | 0.644 | 0.810 | -7.123 |
| TME | PD-1 | Melanoma | -0.995 | 0.601 | 0.852 | -0.603 |
| TME | PD-1 | Other | 0.483 | 0.667 | 0.988 | 1.381 |
| TME | PD-1 | Sarcoma | -1.450 | 1.377 | 0.974 | -0.950 |
| TME | PD-L1 | Carcinoma | 0.055 | 0.852 | 0.566 | 15.404 |
| TME | PD-L1 | Melanoma | 0.164 | 0.856 | 1.011 | 5.223 |
| TME | PD-L1 | Other | 0.329 | 0.882 | 1.449 | 2.679 |
| TME | PD-L1 | Sarcoma | -1.435 | 0.517 | 0.366 | -0.361 |
| TME | CD45 | Carcinoma | 0.354 | 0.788 | 0.791 | 2.225 |
| TME | CD45 | Melanoma | 0.492 | 0.870 | 1.117 | 1.767 |
| TME | CD45 | Other | 0.883 | 1.100 | 1.903 | 1.246 |
| TME | CD45 | Sarcoma | -1.056 | 0.297 | 0.210 | -0.282 |
| TME | Fibronectin | Carcinoma | 0.896 | 0.604 | 0.859 | 0.674 |
| TME | Fibronectin | Melanoma | -1.209 | 0.697 | 1.228 | -0.577 |
| TME | Fibronectin | Other | 0.536 | 1.272 | 2.097 | 2.375 |
| TME | Fibronectin | Sarcoma | -0.972 | 1.244 | 0.880 | -1.281 |
| TME | Ki-67 | Carcinoma | -0.323 | 0.540 | 0.772 | -1.672 |
| TME | Ki-67 | Melanoma | -0.124 | 0.850 | 1.274 | -6.859 |
| TME | Ki-67 | Other | 0.142 | 1.183 | 0.932 | 8.316 |
| TME | Ki-67 | Sarcoma | -0.591 | 0.390 | 0.276 | -0.661 |
| TME | HLA-DR | Carcinoma | 0.188 | 0.707 | 0.846 | 3.763 |
| TME | HLA-DR | Melanoma | 0.178 | 0.992 | 1.179 | 5.558 |
| TME | HLA-DR | Other | 0.929 | 1.061 | 1.282 | 1.142 |
| TME | HLA-DR | Sarcoma | -0.964 | 0.238 | 0.168 | -0.247 |

**Supplementary Table 6.** Performance of ROI selection at a manually predefined input grid. Performance metrics aggregated at the tumor type level and stratified for training and test sets. SD: standard deviation, 95%CI: 95% confidence interval, mAP0.5: mean Average Precision at 0.5 intersection-over-union, mAP0.5-0.95: mean Average Precision at 0.5-0.95 intersection-over-union.

| Performance metric | | Carcinoma | Melanoma | Sarcoma | Other |
|---|---|---|---|---|---|
| Training and Validation set | | | | | |
| Number of slides | | 35 | 6 | 16 | 1 |
| Precision | Mean | 0.967 | 0.973 | 0.965 | 0.984 |
| | SD | 0.011 | 0.006 | 0.003 | – |
| | 95%CI | 0.963 – 0.971 | 0.957 – 0.987 | 0.958 – 0.972 | – |
| Recall | Mean | 0.989 | 0.986 | 0.991 | 0.993 |
| | SD | 0.006 | 0.007 | 0.007 | – |
| | 95%CI | 0.988 – 0.992 | 0.979 – 0.993 | 0.988 – 0.995 | – |
| mAP0.5 | Mean | 0.990 | 0.993 | 0.991 | 0.997 |
| | SD | 0.006 | 0.007 | 0.006 | – |
| | 95%CI | 0.988 – 0.992 | 0.986 – 1.00 | 0.987 – 0.994 | – |
| mAP0.5-0.95 | Mean | 0.902 | 0.907 | 0.909 | 0.909 |
| | SD | 0.023 | 0.027 | 0.024 | – |
| | 95%CI | 0.894 – 0.910 | 0.878 – 0.935 | 0.896 – 0.922 | – |
| Test set | | | | | |
| Number of slides | | 21 | 31 | 57 | 10 |
| Precision | Mean | 0.970 | 0.969 | 0.971 | 0.969 |
| | SD | 0.012 | 0.012 | 0.011 | 0.009 |
| | 95%CI | 0.965 – 0.975 | 0.964 – 0.973 | 0.968 – 0.974 | 0.962 – 0.976 |
| Recall | Mean | 0.989 | 0.989 | 0.991 | 0.991 |
| | SD | 0.006 | 0.007 | 0.007 | 0.007 |
| | 95%CI | 0.986 – 0.992 | 0.987 – 0.992 | 0.989 – 0.992 | 0.986 – 0.996 |
| mAP0.5 | Mean | 0.989 | 0.990 | 0.991 | 0.989 |
| | SD | 0.005 | 0.006 | 0.007 | 0.007 |
| | 95%CI | 0.987 – 0.992 | 0.988 – 0.992 | 0.989 – 0.992 | 0.984 – 0.995 |
| mAP0.5-0.95 | Mean | 0.916 | 0.911 | 0.903 | 0.906 |
| | SD | 0.022 | 0.029 | 0.022 | 0.021 |
| | 95%CI | 0.906 – 0.926 | 0.899 – 0.921 | 0.897 – 0.909 | 0.891 – 0.921 |

**Supplementary Table 7** Parameters used in QuPath of nuclear-like structure segmentation used for validation of the co-registration of multiplex immunofluorescence (mIF) to hematoxylin and eosin (H&E) images. After segmentation, binary masks were exported and further processed.

| Parameters | mIF | H&E |
|---|---|---|
| Detection channel | Syto13 (nuclear) | Hematoxylin OD |
| Background radius (um) | 8.0 | 8.0 |
| Sigma (um) | 1.5 | 1.5 |
| Minimum area ($um^2$) | 10.0 | 10.0 |
| Maximum area ($um^2$) | 400.0 | 400.0 |
| Threshold | 100 | 100 |
| Cell expansion (um) | 1.0 | 1.0 |
| Smooth boundaries | True | True |

121 **Annex I: Patient Mosaic Team**

Nadim J Ajami
Azad Ali
Franklin Alvarez
Brittany Alverez
Bianca Amador
Surosh Avandsalehi
Claudia Alvarez Bedoya
Katrice Bogan
Elena Bogantenkova
Elizabeth Bonojo
Maria Neus Bota-Rabassedas
Elizabeth M Burton
Noble Cadle
Vanessa Castro
Chi-Wan Chow
Randy Aaron Chu
Candace Cunningham
Carrie Daniel-MacDougall
Nana Kouangoua Diane C
Mary Domask
Sheila Duncan
Andrew Futreal
Vivian Gabisi
Jessica Gallegos
Andrea Galvan
Ana Garcia
Jose Garcia
Celia Garcia-Prieto
Christopher Gibbons
Jonathan Benjamin Gill
Dominic Guajardo
Curtis Gumbs
Kristin J Hargraves
Tim Heffernan
Joshua Hein
Sharia Hernandez
Charlotte Hillegass
Yasmine M Hoballah
Theresa Honey
Chacha Horombe
Habibul Islam
Stacy Jackson
Jeena Jacob
Akshaya Jadhav
Robert Jenq
Weiguo Jian
Juliet Joy
Isha Khanduri
Walter Kinyua
Laura Klein
Mark Knafl
Larisa Kostousov
Ying-Wei Kuo
Wenhua Lang
Barrett Craig Lawson
Alexander Lazar
Jack Lee
Erma Levy
XiQi 'Cece' Li
Latasha D Little
Yang Liu
Yan Long
Vielka Lopez
Wei Lu
Sandra Lugo
Aaliyah Maldonado
Jared Malke
Asri Margono
Dipen Maheshbhai Maru
Grace Mathew
Brian McKinley
Jennifer Leigh McQuade
Courtney McRuffin
Gertrude Mendoza
Christopher Miller
Raymond Montoya
Francisco Motemayor
Theresa Nguyen
Heather Perez
Juan Posadas Ruiz
Sabitha Prabhakaran
Mallory Psenda
Gabriela Raso
Mike Roth
Pranoti Sahasrobhojane
Amber Savant
Keri L Schadler
Alejandra Serrano
Kenna R Shaw
Julie M Simon
Elizabeth Sirmans
Luisa Maren Solis Soto
Xingzhi 'Henry' Song
Meghan Stennis
Huandong 'Howard' Sun
Maria Chang Swartz
Marialeska Tariba-Edick
Christopher Vellano
Angela Walker
Ignacio Ivan Wistuba
Scott Eric Woodman
DeArtura Young
Jianhua 'John' Zhang
Haifeng Zhu
Hui 'Helen' Zhu
Olga Bat
Shadarra Crosby
Ellie Freebern
Cindy Hwang
Diana Kouangoua
Yang Li
Sharon Miller
Xiaogang 'Sean' W

1
